# Thioredoxin-1 inhibits granulosa cell ferroptosis to rescue ovarian aging through mitophagy-dependent activation of BNIP3L

**DOI:** 10.1101/2025.07.12.664497

**Authors:** Wenting Yang, Xiaorui Zhang, Rui Xu, Zhuoqi Huang, Yanmin Ma, Jiao Yang, Zheng Lin, Gang Ma, Jianfei Xu, Yu Qiao, Zhen Xiao, Jing Cao, Shan Li, Xinyu Zhang, Andrew P. Hutchins, Guoqing Tong

**Author notes:** These authors contributed equally (Wenting Yang, Xiaorui Zhang, Rui Xu, Zhuoqi Huang). These authors jointly supervised this work (Andrew P. Hutchins, Guoqing Tong).

## Abstract

Ovarian aging is closely associated with a decline in fertility and an increase in reproductive dysfunction. Ovarian granulosa cells (GCs) support oocyte homeostasis and development, yet insight into GC dysfunction during aging is limited. Here, we show that aged GCs of humans and mice have indications of elevated ferroptosis, including increased ferroptosis-related metabolites, lipid peroxidation, and iron accumulation. The ferroptosis inhibitor Ferrostatin-1 reversed ovarian impairment and fertility of aged mice *in vivo*. We show that the age-related reduction in the expression of TXN (thioredoxin) leads to ferroptosis in human and mouse GCs by blocking BNIP3L-dependent mitophagy. Exogenous activation of TXN could promote mitophagy, thereby clearing excessive ROS and inhibiting ferroptosis. These results suggest that anti-ferroptosis-related treatments may assist in treating aging-related reproductive disorders.

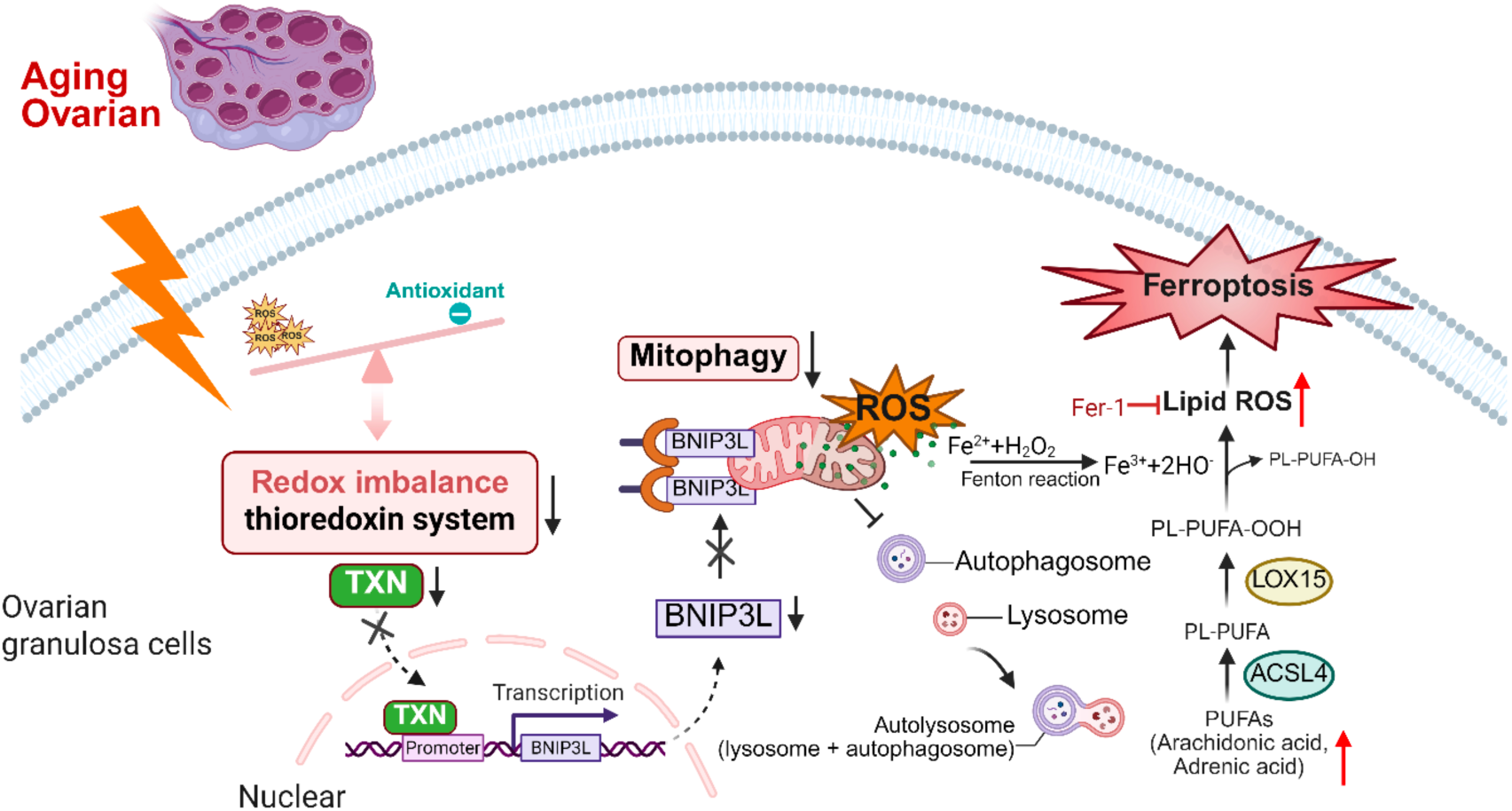

**Key points:**

- Ferroptosis signatures are upregulated in aged GCs from human and mouse ovaries
- TXN is deregulated in aged GCs, leading to mitochondrial and ROS metabolic dysfunction
- TXN binds to DNA to regulate autophagy and mitophagy genes, including *BNIP3L*
- Inhibition of ferroptosis can ameliorate GC dysfunction

## Introduction

Ovarian aging contributes to the progressive decline in female reproductive capability and fertility by decreasing the quality and quantity of oocytes ^1^. Although the contribution of endocrine, genetic, and metabolic factors that cause decreased oocyte quality in ovarian aging has been explored^2^, practical strategies to improve or delay ovarian aging remain limited, and new methods focusing on improving ovarian function are required. In both mice and humans, multiple studies have described the cellular and molecular changes that are associated with ovarian aging ^1,3–6^. Aging transforms the ovary from a youthful, functional state to a senescent one through multiple pathways ^7^. Extensive work has focused on oocyte development quality ^8–10^, but less work has been paid to other cells in the ovary. Particularly important are the granulosa cells (GCs) that protect and communicate with oocytes to manage follicle development, steroidogenesis, and endocrine regulation ^11,12^. Dysfunction of GCs causes follicle atresia, premature ovarian failure, and female subfertility or infertility ^11,12^, and affects oocyte quality ^13–15^. Multiple pathways are deregulated in the aging ovarian GCs, particularly increased inflammation ^16^, DNA damage and oxidation ^17^, lipid peroxidation, and reactive oxygen species (ROS) that can serve as biomarkers for ovarian function ^18^. However, clinical interventions based on these pathways remain limited.

Ferroptosis is a type of programmed cell death caused by iron-dependent accumulation of lipid peroxides and ROS ^19–23^. Evidence suggests that ferroptosis is associated with aging and aging-related pathologies, including increased oxidative stress and cell mortality ^24^. Ferroptosis is potentially related to ovarian aging due to high ROS and abnormal lipid metabolism ^21,25^ which disrupts the normally low ROS levels in oocytes and dormant follicles ^26^. Iron metabolism and ROS are dysregulated in polycystic ovary syndrome (PCOS) ^27,28^, and miR-93-5p drives NF-kB apoptosis and ferroptosis in PCOS GCs ^29^. Premature ovarian insufficiency (POI) can be caused by deregulation of the BNC1-NF2-YAP pathway that induces ferroptosis in oocytes ^30^. Ferroptosis is also widely observed in ovarian cancer ^31,32^. However, a relationship between ferroptosis and redox metabolism in aging and GC dysfunction has not been established.

Mitochondrial dysfunction can have a significant impact on aging GCs by disrupting energy homeostasis and redox balance and impairing steroidogenesis and follicular development. Mitophagy, which is the selective degradation of mitochondria by autophagy, is crucial for maintaining cellular homeostasis and is frequently deregulated in aging ^33^. Mitochondrial deregulation is observed in aged oocytes ^8,34^, and GCs ^35^, suggesting that proper mitophagy is essential for maintaining mitochondrial integrity during follicular development. Spermidine supplementation can improve mitochondrial quality by enhancing mitophagy and likely contributes to improved fertility in aged mice ^36^. However, how mitophagy is mechanistically regulated in the aging ovary remains unclear. Potentially, the accumulation of damaged mitochondria could increase the susceptibility to ferroptosis due to excessive lipid peroxidation and production of ROS ^37,38^. Thioredoxin (TXN) combats ROS-induced oxidative stress and has an inhibitory role in ferroptosis by detoxifying lipid ROS by activating GPX4 (glutathione peroxidase 4) ^39,40^. However, whether TXN functions in ovarian aging and reproductive capability remains unclear. Crosstalk between mitophagy, ROS response (via TXN and GPX4), and ferroptosis could have implications for understanding the complex mechanisms underlying ovarian function decline and age-related pathologies. In this study, we propose a link between ferroptosis, mitophagy, and antioxidant regulation in ovarian aging in GCs. Our results show that human GCs from advanced maternal age women (>40 years) have increased signs of ferroptosis and decreased mitochondrial function. This is accompanied by decreased TXN activity, which leads to impaired BNIP3L-dependent mitophagy, lipid peroxidation, and ROS accumulation that feeds back to cause increased ferroptosis. Inhibition of ferroptosis or activation of TXN improved the quality of aging GCs in mouse and human models and improved fertility in old mice and premature ovarian failure (POF) mouse models.

## Results

### Age-associated senescence and ferroptosis are upregulated in human and mouse ovarian granulosa cells

A lot of attention has been paid to understanding aging in the context of zygote development and implantation ^9,10,41^, however, less effort has been expended on the ovary, particularly on GC support cells ^11^. Hence, we explored signs of senescence that may explain GC dysfunction. Patients with PCOS, premature ovarian insufficiency (POI), and other ovarian hypofunctions caused by tumors or endocrine diseases were excluded from this study. Only patients requiring assisted reproductive treatment (ART) between 22-46 years old were included. GCs were purified from young (<=29 years old), middle-aged (30-36 years old), and older maternal-age patients (>36 years old) by excluding CD45+ mononuclear cells by FACS (**Supplementary Figure 1a**). GC purity was confirmed by immunostaining with FSHR (follicle-stimulating hormone receptor) (**Supplementary Figure 1b**). Interestingly, senescence-associated SA-β-galactosidase (SA-β-gal) and p21^CDKN1A^ were both elevated in GCs from older patients (**Figure 1a-c**). Whilst LMNB1 (Lamin B1), a negative senescence marker, declined with age (**Figure 1b and c**). We reanalyzed scRNA-seq (single-cell RNA-seq) data from human ovaries from young and old patients and isolated the GCs (**Figure 1d and Supplementary Figures 1c**) ^42^. As previously reported, there were three main clusters of GCs: cluster 1 comprised young GCs, cluster 3 was mainly old, and cluster 2 was intermediate (**Supplementary Figure 1d and e**). Aged GCs had elevated levels of ferroptosis-driver genes and decreased expression of ferroptosis-suppressors (**Figure 1d-f and Supplementary Figure 1f, g**) ^43^, and increased senescence genes (**Supplementary Figure 1h**). Interestingly, based on the scRNA-seq, elevated ferroptosis was mainly restricted to GCs and monocytes, and other ovary tissues were relatively unaffected (**Supplementary Figure 1i**). In support of enhanced ferroptosis in GCs, the ferroptosis-related metabolite MDA (Malondialdehyde), an end-product of lipid peroxidation ^22^, was significantly increased from young to old GCs (**Figure 1g**). Conversely, GSH (reduced glutathione), which is the substrate required by GPX4 to reduce lipid peroxides and protect against ferroptosis ^40,44^, was significantly decreased, suggesting GPX4 activity is impaired (**Figure 1h**). Additionally, old GCs had increased ferrous (Fe^2+^) ion deposition, as measured by FerroOrange (**Figure 1i**). Further, suggestive of ferroptosis, mitochondrial membrane potential was reduced in old GCs (**Figure 1j**). These data support elevated ferroptosis-related pathways and proteins in GCs of older maternal-age women.

**Figure 1.**
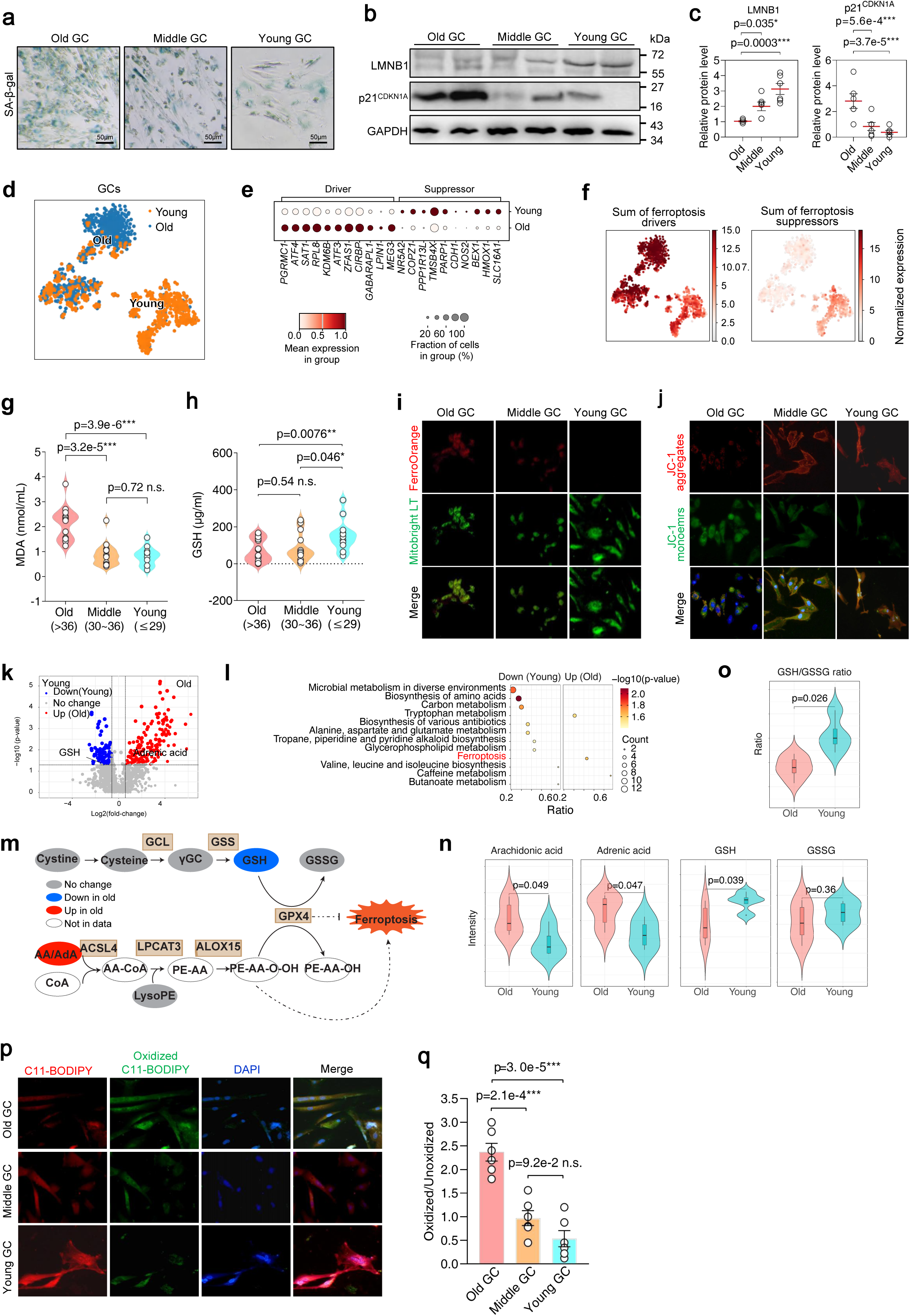
Aging induces senescence and ferroptosis in human ovarian granulosa cells. **a** Representative images of senescence-associated β-galactosidase (SA-β-gal) staining in GCs (Granulosa cells) from old (n=6, >36 years), middle-aged (n=6, 30-36), or young (n=6, <30) women. **b** Western blot of LMNB1 (Lamin B1), p21^CDKN1A^, and GAPDH in GCs from old, middle-aged, and young women. The experiment was performed six times with similar results. **c** Quantitation of the protein levels of LMNB1 and p21^CDNK1A^ based on western blots relative to GAPDH. The p-value is from a one-way analysis of variance (ANOVA) and Tukey’s HSD (Honestly Significant Difference) test (n=6). **d** tSNE (t-stochastic neighbor estimation) embedding of scRNA-seq data from young and old GCs. Cells are colored by the young and old classification, as in the original study. Data is from a reanalysis of Ref. ^42^. **e** Bubble plot showing the mean expression in the scRNA-seq data for a selection of ferroptosis-driver and suppressor genes. **f** tSNE plots of scRNA-seq data (as in **panel d**) colored by the sum of expression of ferroptosis-drivers and suppressors, as defined in Ref. ^43^. **g** Violin plot of the concentration of the end-product of lipid peroxidation, malondialdehyde (MDA), in GCs from old (n=30), middle (n=28), and young (n=22) women. The error bars and statistical analysis were measured by one-way ANOVA and Tukey’s HSD test. **h** The levels of glutathione (GSH) were measured by colorimetric assay for GCs from old (n=30), middle (n=28), and young (n=22) women. The error bars and statistical analysis were measured by one-way ANOVA and Tukey’s HSD test. **i** Microscope images comparing ferrous ion (Fe^2+^; FerroOrange) levels and active mitochondria (Mitobright LT) in old, middle, and young patients. Experiments were performed three times with similar results. **j** Microscope images of mitochondrial membrane potential (MMP) as detected via JC-1 monomers (low MMP; green) and aggregates (high MMP; red) in GCs from old, middle-aged, and young women. Experiments were performed three times with similar results. **k** Volcano plot of metabolites in young and old GCs. Significance was determined as an absolute fold-change > 2 and a Bonferroni-Hochberg adjusted p-value of < 0.05. **l** Ontology analysis of the significantly up- and down-regulated metabolite-related terms. **m** Schematic map for Ferroptosis based on KEGG Ferroptosis pathway (hsa04216). Metabolites are in circles; proteins are in square boxes. **n** Violin plots showing quantitation of arachidonic acid, adrenic acid, GSH, and GSSG for young and old GCs. **o** Violin plot showing the ratio of GSG/GSSG metabolites from the mass spec data. **p** Microscope images of lipid peroxide levels in GCs as measured by C11-BODIPY 581/591 fluorescent probes for the peroxidized (green) and total (red) lipids. The experiment was performed three times with similar results. **q** Quantification of the ratio of lipid peroxidation levels by FACS in GCs from old, middle, and young GCs measured using C11-BODIPY. The error bars indicate the mean ± standard deviation (SD). Statistics are from a two-sided Kruskal-Wallis test.

To explore ferroptosis-related metabolic changes in more detail ^45^, we performed metabolomic mass spectrometry on purified human GCs from young and old women (**Supplementary Figure 2a**). Overall, 1198 metabolites were detected, with 102 metabolites downregulated in young and 146 metabolites up-regulated in old GCs (**Figure 1k, Supplementary Figure 2a-c, and Supplementary Table 1**). Metabolic Ontology analysis indicated the up-regulation of ferroptosis-related metabolites in old GCs (**Figure 1l and m**), particularly GSH, and the downregulation of arachidonic acid (AA; **Figure 1l-n**). The ratio of GSH/GSSG was lower in old GCs (**Figure 1o**), supporting impaired GPX4 catalytic activity. Lipid peroxidation is both a driver and marker for ferroptosis ^46^, hence we used BODIPY-C11 staining to determine the level of lipid peroxidation, which was indeed up-regulated in the old GCs (**Figure 1p and q**). These results show that human GCs from old patients have increased markers of senescence and ferroptosis.

To explore ferroptosis in aged GCs, we utilized the human ovarian GC KGN cell line and treated the cells with increasing concentrations of H_2_O_2_. This led to a decrease in LMNB1 levels, and a corresponding increase in the senescence markers p16^CDKN2A^ and p21^CDKN1A^ were matched by increases in SA-β-gal (**Supplementary Figure 3a and b**), mimicking the phenotype in primary human GCs. As expected, cell viability was significantly reduced in the H_2_O_2_-treated cells compared to the control group (**Supplementary Figure 3c**). The cell viability could be partially rescued when KGN cells were treated with the ferroptosis inhibitor Fer-1 (Ferrostatin-1) (**Supplementary Figure 3c**). Fer-1 treatment also led to a decrease in the ferroptosis metabolite MDA and an increase in GSH (**Supplementary Figures 3d and e**). Lipid ROS levels were significantly increased in H_2_O_2_-treated KGN cells compared with control cells, and this effect could be reversed by treatment with Fer-1 (**Supplementary Figure 3f and g**). Ferrous (Fe^2+^) ion levels were increased, and mitochondrial membrane potential was decreased in H_2_O_2_-treated KGN cells compared with the control (**Supplementary Figure 3h-j**). Together, increased lipid ROS and MDA, and decreased GSH and mitochondrial membrane potential in H_2_O_2_-treated KGN cells suggest ferroptosis is induced in aged GCs and may be related to the dysfunction of ovarian GCs.

Aging impairs reproductive capacity in mice and leads to reduced litter size and disrupted estrous cycles. GCs from old (52 weeks) mice, as in humans, showed a similar pattern of aging-related senescence markers, including increased p21^CDNK1A^ and decreased LMNB1 in old mice (**Supplementary Figure 4a**). The ovarian follicle counts of old mice were significantly lower than young mice (8 weeks) (**Supplementary Figure 4b**), and the number of oocytes was significantly reduced in old mice (**Supplementary Figure 4c**). Furthermore, ovarian tissues showed increased MDA and decreased GSH that correlated with the increasing age of the mice, especially after 48 weeks (**Supplementary Figure 4d and e**). Fe^2+^ was increased, and mitochondrial membrane potential was decreased in GCs of old mice compared with young mice (**Supplementary Figure 4f and g**), suggesting that GCs from old mice also have increased levels of ferroptosis.

### Inhibition of ferroptosis improves ovarian function and fertility in old mice

To define the damage caused by ferroptosis to female fertility *in vivo*, we built a ferroptosis inhibition and activation mouse model by repeated intraperitoneal injection of the ferroptosis inhibitor Fer-1 in old mice, and ferroptosis initiator Erastin in young mice (**Figure 2a**). We also generated a premature ovarian failure (POF) mouse model induced by busulfan and cyclophosphamide treatment (**Figure 2a**) ^47^. Ovarian size weight (ovarian weight/body weight) of 12-week-old mice treated with Erastin was decreased in the POF mice (**Figure 2b and c**). Notably, ovarian weight was rescued by Fer-1 injection in both 52-week-old and POF mice (**Figure 2b and c**). All follicle types were significantly decreased in the Erastin-treated and POF groups, and several follicle types were significantly up-regulated in the old or POF mice treated with Fer-1 (**Figure 2d**). This effect extended to litter size, as the average litter size was significantly smaller in both Erastin-treated and POF mice compared to their respective controls (**Figure 2e and f**), and, remarkably, 52-week-old mice treated with Fer-1 gave birth (**Figure 2e and f**). Erastin-treated and old mice had low hormone levels that could be improved with Fer-1 treatment (**Figure 2g**). Fer-1 and Erastin cause systemic effects; hence, to remove potential confounding effect of the mouse endometrium on implantation rates, we performed *in vitro* fertilization of POF and aged mice oocytes and monitored blastocyst development *in vitro*. Quantification of cleavage rates showed a significant increase in the 2-cell (2C) cleavage rate in embryos from old mice treated with Fer-1, and a significant reduction in 2C-cleavage in young mice that had been treated with Erastin (**Figure 2h and i**). Fer-1 also significantly improved 2C-cleavage in the POF model (**Figure 2h and i**). This shows that inhibition of ferroptosis in mice improved oocyte quality and embryonic developmental potential.

**Figure 2.**
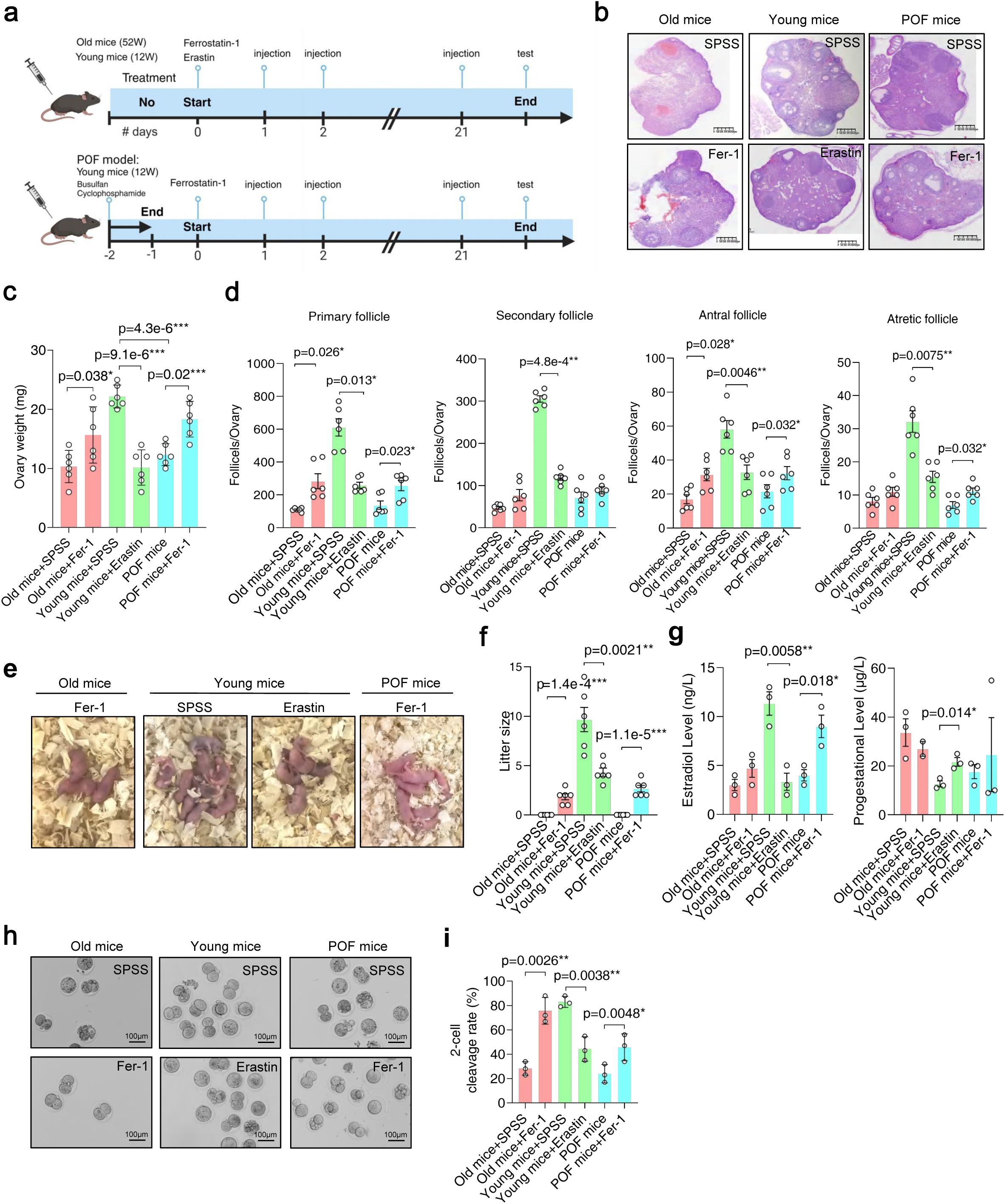
Increased ferroptosis in GCs of aged and premature ovarian failure model mice. **a** Schematic of the mouse models used in this study. Young (<12 weeks) or old (>52 weeks) mice were treated with Fer-1 (Ferroptosis inhibitor; 1 mg/kg) or Erastin (20 mg/kg; Ferroptosis inducer) via intraperitoneal injection once a day for 21 days. To induce POF (premature ovarian failure), 12-week-old mice were administered a single dose of busulfan and cyclophosphamide. Six mice were used for each group. **b** Hematoxylin and eosin-stained ovary sections from 12- or 52-week-old mice treated with SPSS (vehicle), Fer-1, Erastin, or BuCy. Scale bar = 625 μm. **c** Ovary weight of 12- and 52-week-old mice treated with Erastin, or in POF model mice (n=6). Statistics are from an unpaired two-tailed Student’s t-test. **d** Bar chart of the number of follicles per ovary for the indicated mice and treatments for primary, secondary, antral, and atretic follicles (n=6 per group). Statistics are from an unpaired two-tailed Student’s t-test. **e** Example images of litter size from 12-week and 52-week mice treated with the indicated chemicals. 52-week-old SPSS (vehicle) and 12-week-old BuCy-treated mice did not produce offspring. **f** Bar chart of average litter sizes of Erastin, BuCy, 12-week-old (young) POF model mice treated with Fer-1 or SPSS (vehicle), and 52-week-old mice treated with Fer-1 (n=6 per group). Statistics are from an unpaired two-tailed Student’s t-test. **g** Bar chart of the levels of estradiol and progesterone for mice with the indicated age and treatments (n=6 per group). Statistics are from an unpaired two-tailed Student’s t-test. **h** Images of 2-cell stage embryos collected from 12 and 52 week-old mice treated with SPSS (vehicle), Fer-1, Erastin, or POF model mice treated with Fer-1. **i** Bar chart showing the 2-cell cleavage rate (n=6) for embryos from the indicated age mice and treatments. Statistics are from a two-sided Kruskal-Wallis test.

### The TXN antioxidant system is deregulated in aged GCs

The data from the mouse models suggest that a consequence of increased ferroptosis is a depleted number of follicles and ovarian function. Hence, we reanalyzed an RNA-seq dataset ^4^ of GCs from six patients with diminished (DOR) or normal ovarian reserves (NOR). In the DOR GCs, 2484 genes were significantly upregulated and 2309 downregulated (**Supplementary Figure 5a**), and as expected, ferroptosis drivers tended to be downregulated in DOR, and suppressors were associated with NOR patients (**Figure 3a and Supplementary Figure 5b, c**). The above data suggested ferroptosis is up-regulated in DOR and may be a contributing factor, and human DOR at least superficially resembles the erastin-treated young mice.

**Figure 3.**
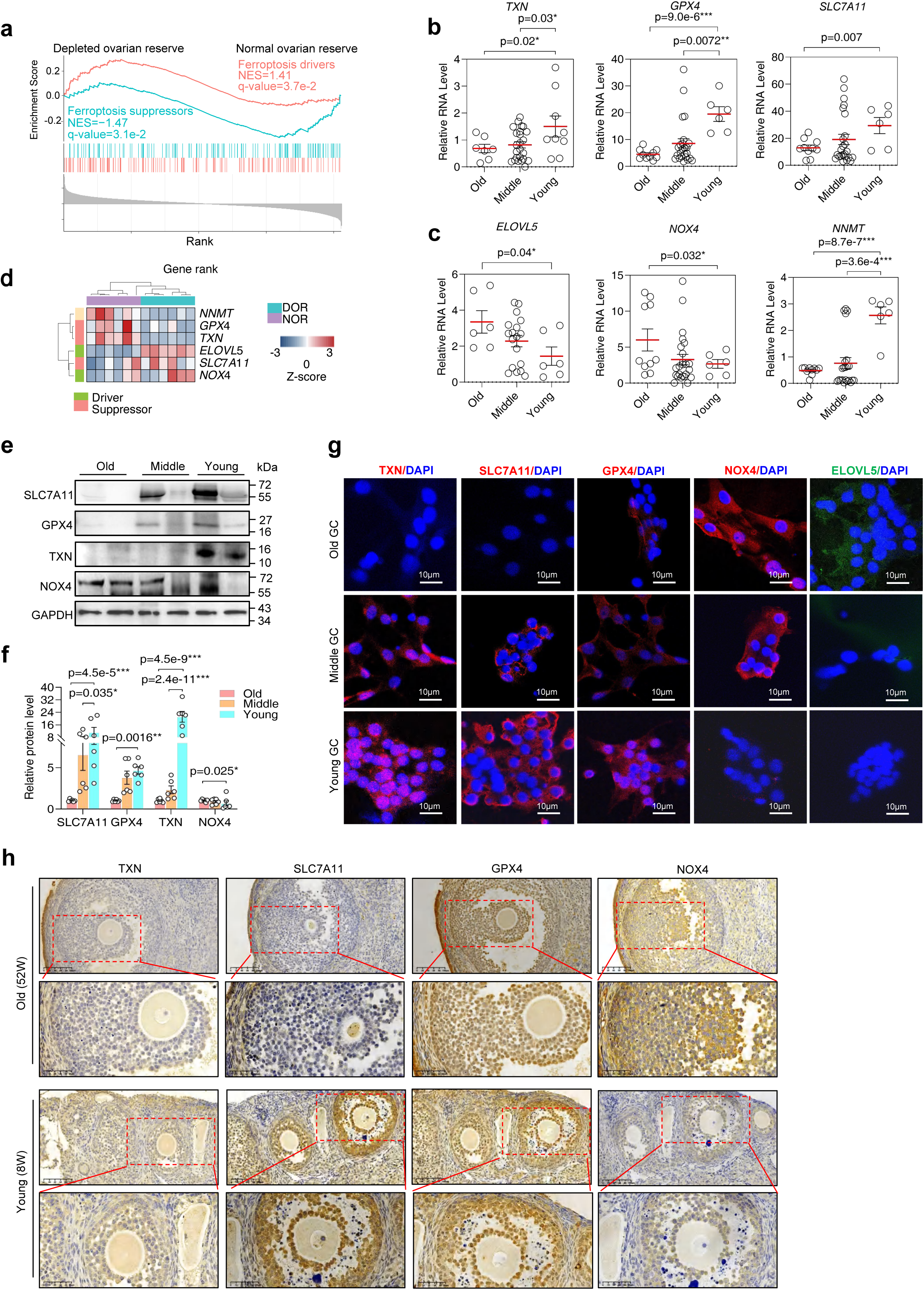
The TXN antioxidant system is impaired in GCs of aged human and mouse ovaries. **a** GSEA for drivers and suppressors for old women (n=6) versus young women (n=6). Data is from a reanalysis of GSE232306 ^4^. Ferroptosis driver or suppressor genes, as defined in FerrDB ^43^. **b** RT-qPCR for the ferroptosis suppressors *TXN*, *GPX4*, and *SLC7A11.* GCs were obtained from old women (n=10, >36 years), middle-aged women (n=18, 30~36), and young women (n=6, <30). The error bars indicate the mean±SD. Statistics are from an independent samples two-sided Student’s *t*-test. **c** RT-qPCR for the ferroptosis drivers *ELOVL5*, *NOX4*, and the ferroptosis activator *NNMT* in GCs. The error bars indicate the mean±SD. Statistics are from an independent samples two-sided Student’s *t*-test. **d** Heatmap of selected differentially regulated ferroptosis drivers and suppressors in DOR (depleted ovarian reservoir) and NOR (normal ovarian reservoir) human samples. Data is from a reanalysis of GSE232306 ^4^. Ferroptosis driver or suppressor genes, as defined in FerrDB ^43^, and the ferroptosis activator *NNMT*. **e** Western blot of SLC7A11, GPX4, TXN, NOX4, and GAPDH in GCs from old, middle, and young women. **f** Bar chart of quantitated levels of SLC7A11, GPX4, TXN, and NOX4 in GCs from old, medium, and young women, relative to GAPDH and relative to the old GC samples. The experiment was performed 6 times with similar results. **g** Immunofluorescence of TXN, SLC7A11, GPX4, NOX4 and ELOVL5, in GCs from old, middle, and young women (n=3 per group). **h** Immunohistochemistry of TXN, SLC7A11, GPX4, and NOX4 in sections of mouse ovaries at 52 weeks old and 8 weeks old (n=3 per group).

Cystine is imported into the cell via the SLC3A2/SLC7A11 (System X_c_^-^), a glutamate/cystine transporter. Cystine is then reduced to cysteine by TXN to modulate GPX4 activity ^48,49^. Erastin promotes ferroptosis by inhibiting SLC3A2/SLC7A11 transportation, suggesting that this process may be disrupted in DOR and aged human GCs. We confirmed by RT-qPCR that the ferroptosis suppressor genes (*TXN*, *GPX4*, and *SLC7A11*) were down-regulated (**Figure 3b**), whilst the ferroptosis drivers *ELOVL5*, *NOX4*, and the ferroptosis activator *NNMT* ^50^ were up-regulated (**Figure 3c**) in GCs from young, middle, and aged human GCs. This pattern was matched in RNA-seq data from NOR and DOR GCs (**Figure 3d**). Interestingly, SLC7A11, GPX4, and TXN were downregulated at the protein level in old GCs from humans, whilst NOX4 was up-regulated in old GCs (**Figure 3e and f**). Immunofluorescence imaging agreed as SLC7A11, TXN, GPX4 were downregulated in the old GCs, whilst ELOVL5 and NOX4 were up-regulated (**Figure 3g**). There was a similar pattern in the ovaries of aged mice, as IHC confirmed that TXN and SLC7A11 were downregulated in 52-week-old mice, whilst NOX4 was up-regulated (**Figure 3h**). GPX4 expression was somewhat mixed in this mouse model experimental system (**Figure 3h**). IHC of human GCs, however, showed that GPX4, TXN, and SLC7A11 were all reduced in GCs from old patients, whilst NOX4 was up-regulated (**Figure 3g**). These data suggest that dysfunction of TXN, GPX4, SLC7A11, and NOX4 may induce or modulate ferroptosis in aged human GCs.

### TXN delays H_2_O_2_-induced senescence in KGN GC-like cells by inhibiting ferroptosis

To address whether deficiency of the TXN-based antioxidant system triggers ferroptosis in aged GCs, we knocked down *TXN* using shRNAs (sh*TXN*#1, sh*TXN*#2) or overexpressed (*TXN* OE#1, *TXN* OE#2) in KGN cells (**Figure 4a**). Overexpression of *TXN* significantly increased cell viability and rescued cell proliferation inhibition in H_2_O_2_-treated KGN cells (**Supplementary Figure 6a**). Notably, and indicative of ferroptosis, Fe^2+^ accumulation (**Figure 4b**) and peroxidized lipids (**Figure 4c and Supplementary Figure 6b**) were significantly increased in *TXN* knockdown cells compared with the sh*LUC* controls. In contrast, *TXN* overexpression resulted in the opposite effect: Fe^2+^ accumulation and lipid peroxidation were low (**Figure 4c and Supplementary Figure 6b**). Notably, in cells stressed with H_2_O_2,_ overexpression of *TXN* reduced lipid peroxidation and Fe^2+^ levels (**Figure 4b, c, and Supplementary Figure 6b**). Notably, mitochondrial depolarization was reduced in *TXN* OE cells and increased in *TXN* knockdown cells (**Supplementary Figure 6c**), suggesting altered mitochondrial function. In addition, SA-β-gal production was reduced in *TXN* OE H_2_O_2_-treated cells but was increased in the *TXN* knockdown cells (**Figure 4d**). In support for a role for TXN in ferroptosis and senescence, the senescence markers (p21^CDKN1A^, p16^CDKN2A^, and LMNB1) and ferroptosis-related markers (GPX4, SCL7A11, and NOX4) were increased in *TXN* knockdown cells and matched the effect of H_2_O_2_ (**Figure 4e**). Overall, TXN levels inversely correlated with higher levels of Fe^2+^ (**Figure 4b**), peroxidized lipids (**Figure 4c**), SA-β-gal (**Figure 4d**), and ferroptosis marker gene expression (**Figure 4e**).

**Figure 4.**
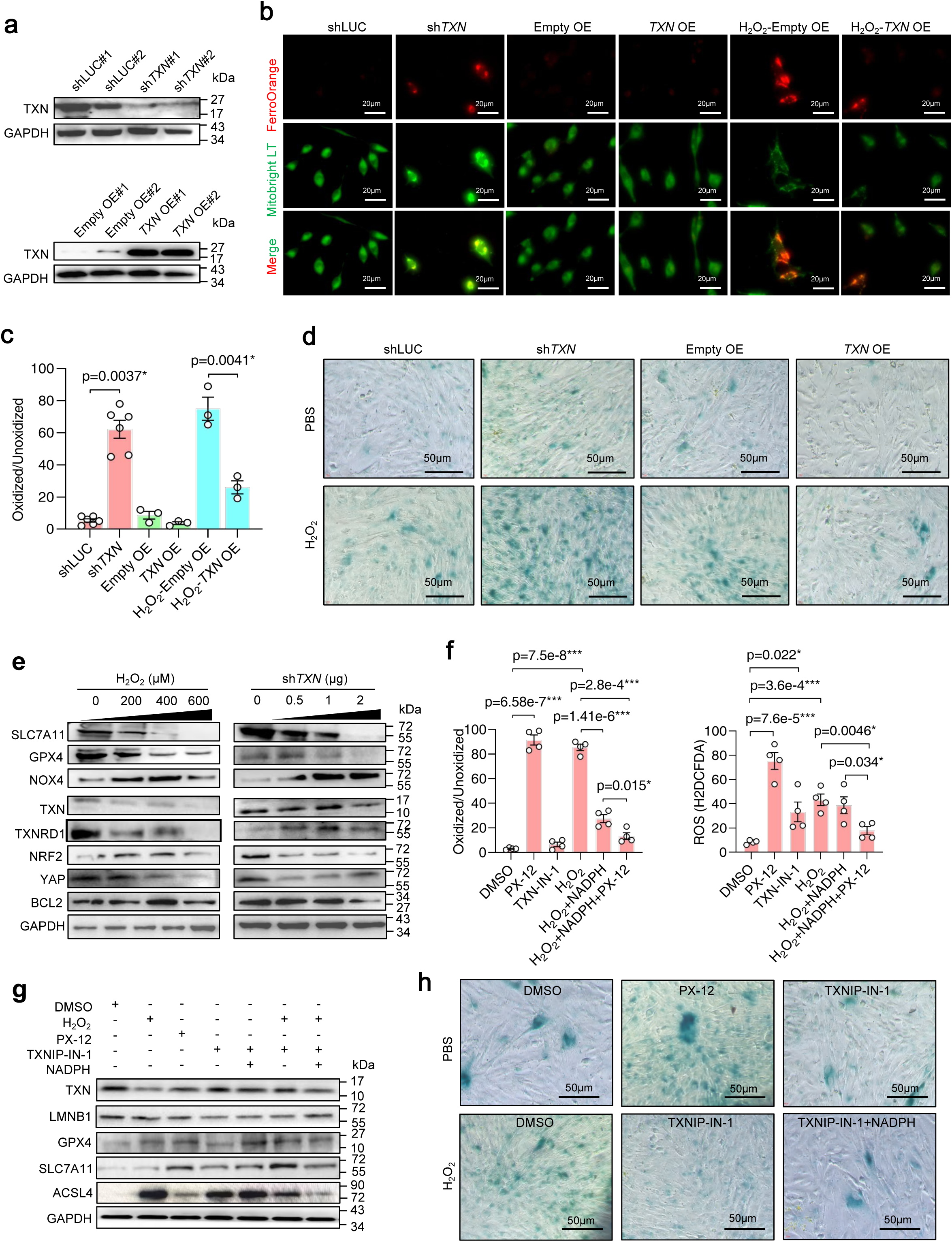
TXN delays the aging phenotype in ovarian GCs by inhibiting ferroptosis. **a** Western blot of TXN protein levels in KGN cells transfected with shRNAs against *TXN* (sh*TXN*#1 and sh*TXN*#2) or overexpressed (*TXN* OE#1 and *TXN* OE#2), and their appropriate controls. **b** Comparison of the ferrous ion (Fe^2+^) level in *TXN* knockdown and overexpressing KGN cells by the Fe^2+^ indicator FerroOrange fluorescent probe (red), and MitoBright LT Green. **c** Lipid peroxidation levels were measured with the C11-BODIPY 581/591 dye by FACS (green/red for oxidized/reduced lipids) in KGN cells transfected with an shRNA targeting TXN or LUC as a control, or a plasmid containing a *TXN* overexpression construct. Cells were also treated with H_2_O_2_ or PBS. The error bars indicate the mean±SD, and the p-value is from a two-sided Kruskal-Wallis test. **d** SA-β-gal staining in TXN-silenced and overexpressed KGN cells treated with H_2_O_2_ or PBS. Scale bar = 50 μm. **e** Western blots for the indicated proteins in KGN cells treated with increasing concentrations of H_2_O_2_ or transfected with increasing amounts of an shRNA targeting *TXN*. **f** Quantitation of oxidized lipids using C11-BODIPY dye (left plot) and total ROS (reactive oxygen species) (right plot) in KGN cells treated with the indicated factors: PX-12 (TXN inhibitor), TXNIP-IN-1 (TXN activator), NADPH, or H_2_O_2_. The error bars indicate the mean±SD, and the p-value is from a two-sided Kruskal-Wallis test. The experiment was performed in biological quadruplicate. **g** Western blot of the indicated proteins in KGN cells treated with DMSO as a control, H_2_O_2_, PX-12 (TXN inhibitor), TXNIP-IN-1 (TXN activator), or NADPH. **h** SA-β-gal staining in KGN cells treated with DMSO, PX-12, TXN-IP-1, with PBS as a control, or H_2_O_2_. Scale bar = 50 μm.

We next used the TXN activators (TXNIP-IN-1 and NADPH) and an inhibitor (PX-12) to explore the role of TXN in ferroptosis. Cell viability was significantly increased by TXNIP-IN-1 when combined with NADPH, and this pattern extended to cells treated with H_2_O_2_ (**Supplementary Figure 6d**). GSH was decreased in PX-12-treated cells and increased when TXN was activated (TXNIP-IN-1 or NADPH) (**Supplementary Figure 6e**). Conversely, MDA levels were significantly increased after PX-12 treatment and decreased in cells treated with TXNIP-IN-1 and/or NADPH (**Supplementary Figure 6f**). Furthermore, mitochondrial membrane potential, accumulation of Fe^2+^, lipid peroxidation, and ROS levels were all decreased in PX-12-treated cells (**Figure 4f and Supplementary Figure 6g-j**). Notably, these ferroptosis-related factors were significantly increased by PX-12 treatment and reduced by TXNIP-IN-1 or NADPH (**Figure 4f and Supplementary Figure 6g-j**). These changes were reflected in changes in protein levels, as the ferroptosis suppressor proteins SLC7A11 and GPX4 were significantly up-regulated by PX-12 treatment (**Figure 4g**). Conversely, treatment with the TXN activators TXNIP-IN-1 and NADPH led to upregulation of ACSL4 (**Figure 4g**). Finally, SA-β-gal was increased in H_2_O_2_ and PX-12-treated cells and could be ameliorated by TXN-IN-1 and NADPH (**Figure 4h**). These results suggested that activation of TXN significantly inhibited ferroptosis in H_2_O_2_-treated KGN cells.

### TXN deficiency blocks mitophagy in aged ovarian GCs

To study the mechanism underlying the ferroptosis induced by TXN deficiency in ovarian aging, we performed a 4D-DIA proteomics to determine the differentially abundant proteins in human ovarian GCs from young and old individuals and identified 8127 proteins, of which 71 and 436 were differentially abundant in old versus young patients (**Figure 5a and Supplementary Table 2**). KEGG pathway enrichment analysis showed that, among the top 10 enriched pathways for the young vs old, mitophagy and autophagy-related pathways were significantly enriched in young (**Figure 5b and Supplementary Figure 7a and b**), including mitophagy-related proteins, such as BCL2L1, OPTN, and BNIP3L (**Figure 5c and Supplementary Figure 7b**). Interestingly, *BNIP3L* RNA was modestly downregulated (**Supplementary Figure 7c**) and was significantly downregulated in the mass spec data, western blot, and immunofluorescence of old and young GCs (**Figure 5c-e**).

**Figure 5.**
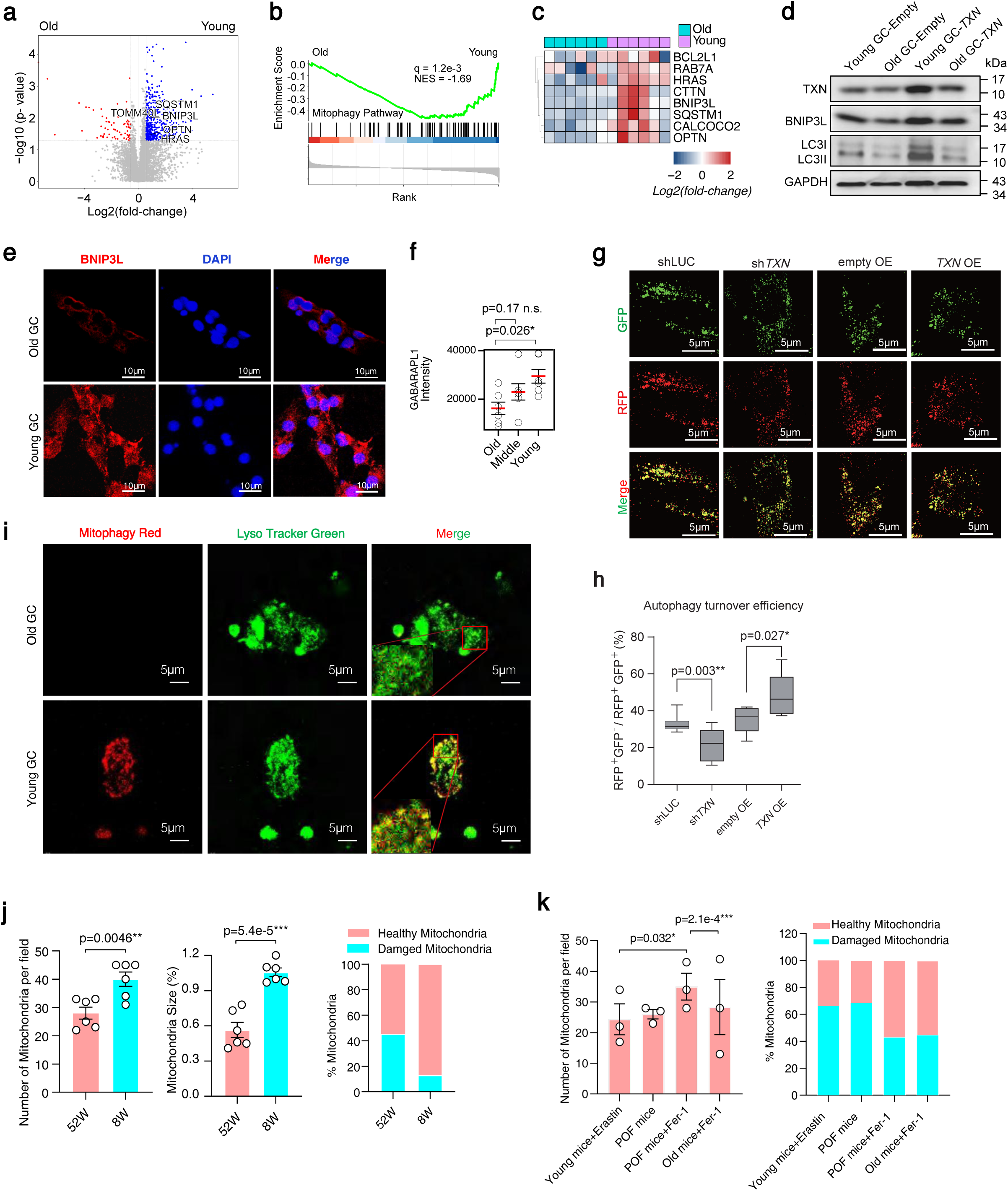
TXN deficiency blocks mitophagy in aged ovarian GCs. **a** Volcano plot of 4D-DIA proteomics for human ovarian GCs from young women (<29 years old, n=6) and old women (>36 years old, n=6). A protein was considered differentially abundant if its p-value was <0.05 and the absolute fold-change > 1.5. **b** GSEA for genes in the mitophagy pathway in old and young individuals. Proteins were ranked by their fold-change from old to young. **c** Heatmap of selected mitophagy-related proteins. **d** Western blot for TXN, BNIP3L, LC31/II and GAPDH as a control for Young and Old GCs, and GCs transfected with a plasmid overexpressing *TXN*. **e** Immunofluorescence of BNIP3L in GCs from old and young women. **f** Dot plot of the mass spec intensity of BNIP3L in young, middle, and aged GCs. The error bars indicate the mean±SD, and the p-value is from a two-sided Mann-Whitney U test **g** Images of autophagic activity based on a GFP-RFP-LC3B reporter. Green puncta indicate nascent autophagosomes, yellow indicates autophagosomes, and red puncta mark autolysosomes. The experiment used KGN cells transfected with an shRNA targeting *LUC* or *TXN*, or an Empty or *TXN* overexpressing vector. **h** Quantification of autophagic flux across different conditions (as in **panel g**), represented by the number of colocalized RFP/GFP voxels per cell. Statistical significance was assessed using a one-sided unpaired Student’s t-test. **i** Fluorescence imaging of mitophagy (Mitophagy Red; becomes red when pH drops in mitochondria) and LysoTracker (lysosomes) in GCs from old and young women. **j** The number of mitochondria (left plot), size (middle plot) and percentage of healthy/damage mitochondria (right plot) as estimated from transmission electron microscopy (TEM) in mice ovary GCs from young mice (8W) and old mice (52W) (See also **Supplementary Figure 7f and g**). The error bars indicate the mean±SD (n=6 for each group). Statistics are from an unpaired two-sided Student’s t-test. **k** Dot plot of the number of mitochondria (left plot), and bar chart of the percentage of healthy/damage mitochondria (right plot), as estimated from TEM, in mice ovary GCs from young mice (8W) treated with Erastin, POF model mice treated with Fer-1 or SPSS and old mice (52W) treated with Fer-1. The error bars indicate the mean±SD (n=3 for each group). Statistics are from an unpaired two-sided Student’s t-test.

LC3/GABARAP proteins are important for autophagy ^51^ (MAP1LC3B) was not detected in the protein mass spec data, however the autophagy LC3B/ATG8-related protein GABARAPL1 was significantly down-regulated at the RNA, and protein level in old GCs (**Figure 5d, f and Supplementary Figure 7d and e**), suggesting impaired autophagy and/or mitophagy. To reveal autophagic flux, KGN cells were transfected with a dual reporter plasmid containing GFP and RFP fused to LC3B (GFP-RFP-LC3B) ^52^. Knockdown of *TXN* in KGN cells led to increased RFP and autophagic turnover, conversely, overexpression of *TXN* led to increased autophagic flux (**Figure 5g and h**), suggesting TXN is driving increased autophagy. To observe mitophagy, we examined the formation of mitophagosomes represented by co-localizing mitochondria and lysosomes ^45^. Indeed, in young GCs, there was a substantial increase in mitophagy and co-localization with lysosomes, and this pattern was absent in GCs from old patients (**Figure 5i**). Similarly, when *TXN* was transfected into young and old patient GCs, young GCs responded with increased LC3II, and presumably autophagy, whilst old cells were weak to respond (**Figure 5d**). These results provide evidence for TXN-driven impaired autophagy and mitophagy in old GCs, due to disrupted TXN regulation.

We explored mitophagy in aged mice ovarian tissues. Transmission electron microscopy (TEM) of mitochondrial membranes suggested dysfunction in mitochondrial membranes as they appeared fuzzy or broken (**Supplementary Figure 7f**). The total number, mitochondrial size, and healthy/unhealthy ratio were all significantly decreased in mouse GCs in 52-week-old mice compared to young 8-week mice (**Figure 5j**). Inhibition of ferroptosis via intraperitoneal injection of the ferroptosis inhibitor Fer-1 improved mitochondrial morphology of GCs (**Supplementary Figure 7g**). Notably, mitochondrial dysfunction, swelling, and membrane potential collapse were all present in Erastin and POF model mice and were rescued with the addition of Fer-1 (**Figure 5k and Supplementary Figure 7h**). The aging-related marker p21^CDKN1A^ was decreased in Fer-1-treated old and POF model mice, and LMNB1 was increased (**Supplementary Figure 7h**). Finally, the ROS markers TXN, NOX4, and ferroptosis markers SLC7A11, GPX4, and ACSL4 were all increased (**Supplementary Figure 7h**), and NAD dehydrogenase-related proteins were also higher in old GCs (**Supplementary Figure 7i**). These data suggest a link between ROS, mitochondrial dysfunction, and ferroptosis.

### TXN regulates the expression of *BNIP3L* to trigger mitophagy in GCs

The mitochondrial outer membrane protein BNIP3L serves as a mitophagy receptor by recognizing autophagosomes through ATG8 ^53^. It was up-regulated in old GCs in the human GC mass spec (**Figure 6a**), suggesting the involvement of BNIP3L-mediated mitophagy. We wondered if *BNIP3L* expression was downstream of TXN. When *TXN* was knocked down or overexpressed in KGN cells BNIP3L protein and RNA matched the TXN changes (**Figure 6b, c and Supplementary Figure 8a and b**), additionally, there was a shift in LC3 from LC3I (cytoplasmic) to the LC3II (autophagosome, membrane-bound) form that signifies increased autophagy, and LC3II/LC3I ratio was reduced in KGN cells treated with H_2_O_2_ and could be rescued when *TXN* was overexpressed (**Figure 6b and c**). This suggests that mitophagy enhanced by TXN might be regulated in a BNIP3L-dependent manner.

**Figure 6.**
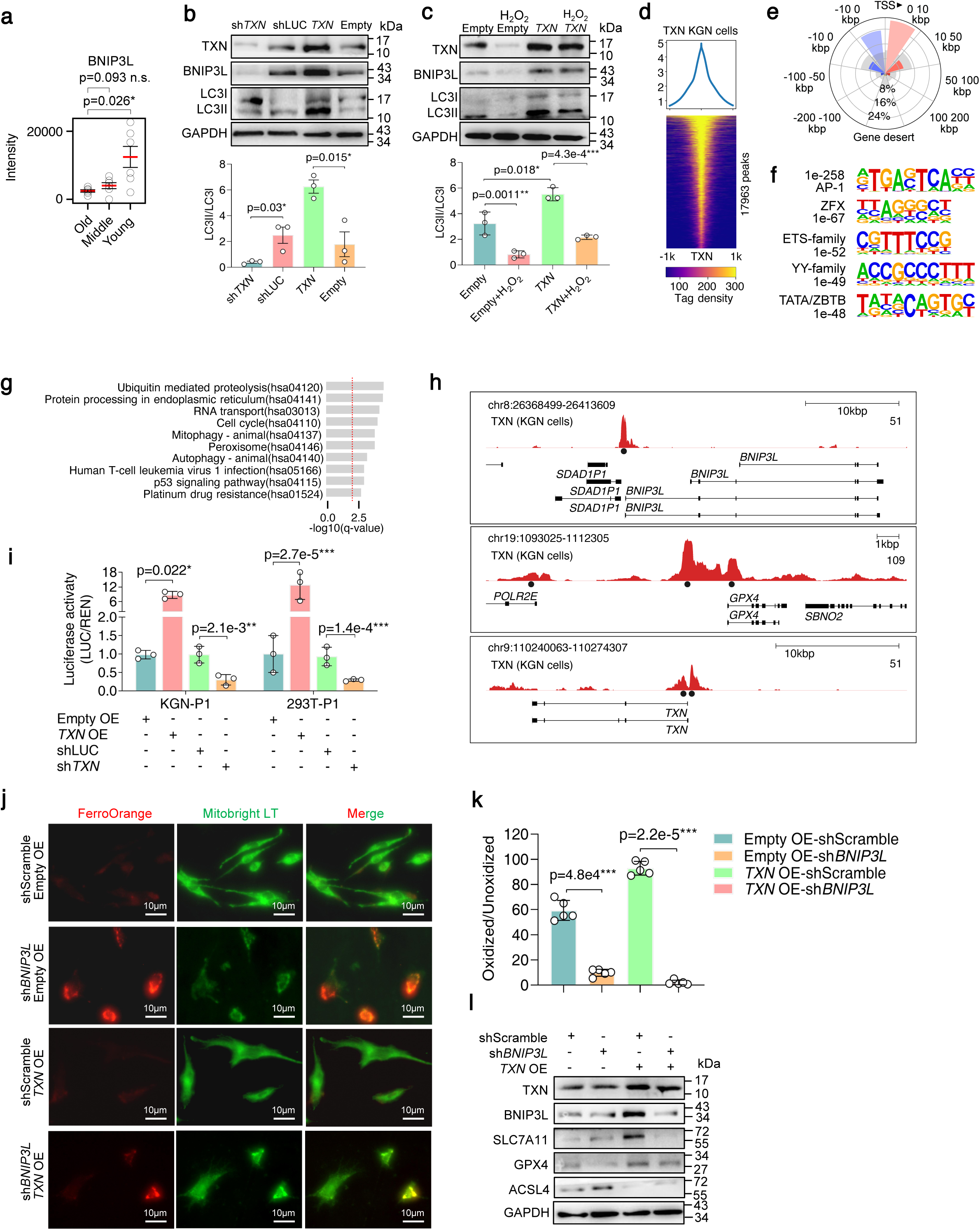
TXN regulates the expression of *BNIP3L* and subsequent ferroptosis in GCs. **a** Protein level of GABARAPL1 in GCs from old, middle-aged, and young human GCs from mass spec data. Significance is from a two-sided Mann-Whitney U test. **b** Western blot for TXN, BNIP3L, and LC3I/II in TXN-knockdown and overexpressing KGN cells, and the relative expression levels relative to GAPDH (bar chart below). **c** Western blot for TXN, BNIP3L, and LC3I/II in TXN overexpressing KGN cells treated with H_2_O_2_ and analysis of relative protein levels relative to GAPDH (bar chart below). **d** Heatmap and pileup of TXN binding to DNA in KGN cells. **e** Genome distribution of the TXN CUT&Tag peaks. Peaks were annotated to the nearest TSS and allocated to bins either 5’ (negative numbers) or 3’ relative to the TSS. A random background is shown in grey for comparison. **f** TF motif discovery at TXN-bound loci. The TF family and q-value are indicated on the left. **g** Gene ontology for KEGG pathways for TXN-bound genes. **h** Genome views around the promoters of *BNIP3L*, *GPX4,* and *TXN*. **i** Dual luciferase reporter assay for the BNIP3L promoter region cloned in front of luciferase in TXN-knockdown and overexpressing KGN and 293T cells. Significance is from a two-way ANOVA and Tukey test. **j** Fe^2+^ and lipid ROS levels in *BNIP3L* knockdown and TXN overexpressing KGN cells. **k** Quantitation of oxidized lipids using C11-BODIPY dye in KGN cells transfected with shRNAs against *BNIP3L* or a scrambled control, or a *TXN,* or an Empty overexpression vector. The error bars indicate the mean values±SDs, and the p-value is from an unpaired two-tailed Student’s t-test. **l** Western blot for TXN, BNIP3L, SLC7A11, GPX4, and ACSL4 in KGN cells transfected with an shRNA targeting *BNIP3L* or a scrambled control, or with a *TXN* overexpression vector.

TXN regulates the activity of transcription factors (TFs), for example, p53 ^54^, NFkB ^55^, and AP-1 ^56,57^. Evidence suggests TXN may directly interact with TFs bound to DNA ^54,57,58^, possibly through APE1/REF1 ^55,56^. Hence, we wondered if TXN could bind to the genome, essentially acting as an epigenetic factor to modulate TF activity, and if so, could this identify downstream regulatory targets of TXN. To this end, we performed CUT&Tag for TXN in KGN cells ^59^. Peak discovery identified 17,963 TXN-bound loci, which were primarily clustered around transcription start sites (TSSs) (**Figure 6d and e**). Motif discovery identified AP-1 motifs at the TXN-bound loci (**Figure 6f**), which suggests indirect binding of TXN to the genome. Interestingly, gene ontology analysis of TXN-bound genes showed pathways related to mitophagy, proteolysis, and autophagy (**Figure 6g and Supplementary Figure 8c**), and this was exemplified by TXN binding to the promoters of *BNIP3L*, *GPX4*, *TXN*, and *SLC7A11* (**Figure 6h and Supplementary Figure 8d**).

We confirmed the binding of TXN to the *BNIP3L* promoter using chromatin immunoprecipitation (ChIP-qPCR) (**Supplementary Figure 8e and f**). To test whether TXN binding regulates *BNIP3L*, we generated a luciferase vector containing the *BNIP3L* promoter (**Supplementary Figure 8e**). The luciferase activity in the *TXN*-overexpressing cells was increased, whereas when *TXN* was knocked down, luciferase activity was reduced in both KGN and 293T cells (**Figure 6i**). These data suggested that TXN binds to DNA to regulate mitophagy and autophagy-related genes through AP-1 and binds to the promoter and activates *BINP3L*.

To confirm that the mitophagy protein BNIP3L was involved in ferroptosis, we transfected a vector containing an shRNA targeting *BNIP3L* and a control scrambled shRNA into *TXN* OE KGN cells. Cell viability was significantly impaired when *BNIP3L* was knocked down in *TXN* OE cells (**Supplementary Figure 8g**). The ferroptosis-related metabolite GSH was significantly lower in the *BNIP3L* knockdown KGN cells, whilst MDA was significantly higher (**Supplementary Figure 6h**). Furthermore, Fe^2+^, lipid ROS, and mitochondrial depolarization were all significantly increased in *BNIP3L* knockdown cells and could be partially rescued in the *TXN* OE cells. (**Figure 6j, k, and Supplementary Figures 8i and j**). Similarly, in KGN cells treated with H_2_O_2_, the ferroptosis markers (lipid ROS and MMP levels) were increased when *BNIP3L* was overexpressed (**Supplementary Figure 8k and l**). Finally, western blot showed that the ferroptosis suppressors *SLC7A11* and *GPX4* were significantly down-regulated, while the ferroptosis driver *ACSL4* was up-regulated in *BINP3L* knockdown cells (**Figure 6l**). These results establish a mechanistic link suggesting TXN regulates *BNIP3L* to inhibit ferroptosis in aged ovarian GCs.

Up-regulation of ferroptosis in GCs may have a knock-on effect on the oocytes they surround and support. Interestingly, we observed an accumulation of Fe^2+^ in the cytoplasm of oocytes from old women that was absent in young women (**Figure 7a and b**). This suggests ferroptosis in ovarian GCs may affect oocyte development (or vice versa) to cause iron overload. This also matches the up-regulation of ferroptosis seen in POI ^30^. Notably, these changes cannot be detected from RNA-seq of oocytes, as ferroptosis and mitophagy-related genes are unchanged between young and old oocytes (**Figure 7c**) ^60^. Overall, ferroptosis in GCs leads to impaired oocyte quality, which leads to reduced fertility in older women.

**Figure 7.**
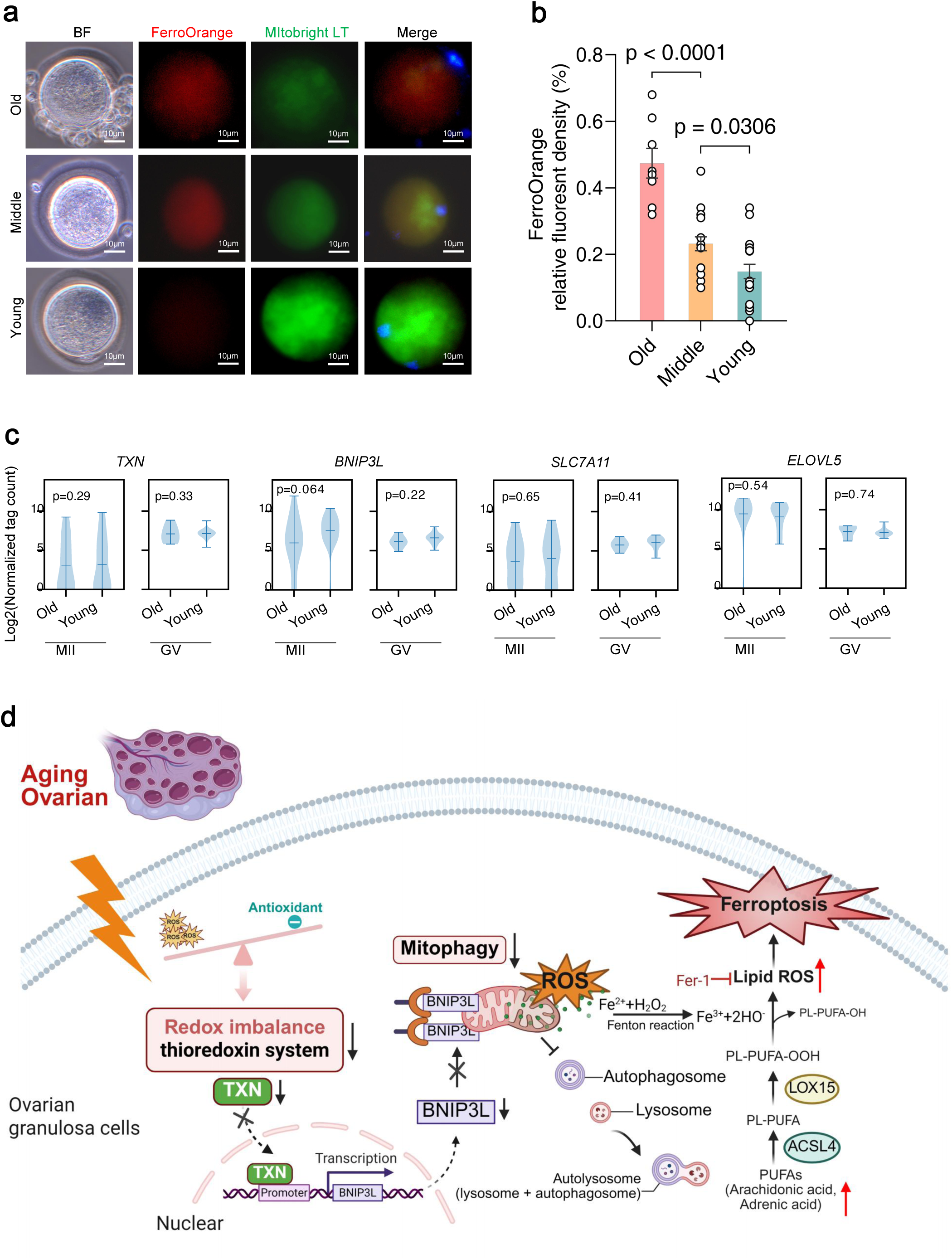
Advanced maternal age oocytes have increased iron accumulation and decreased mitochondria. **a** Co-staining of Fe^2+^ (red) and mitochondria (green) in mature MII oocytes (IVM culture from GV oocytes, 3 independent experiments with a total of >10 oocytes) from old, middle, and young women. **b** Quantitation of FerroOrange fluorescence in MII oocytes from old, middle and young patients. Statistics are from an ANOVA test with Tukey’s HSD. **c** Violin plots of RNA-seq of aged, middle-aged, and young MII oocytes for selected ferroptosis and mitophagy genes. Note that *NOX4*, *GPX4*, and *NNMT* are not expressed/detectable. Data is from a reanalysis of GSE158802 ^60^. Significance is from a two-sided Welch’s t-test. **d** Model of ferroptosis in age-related GC and oocyte degradation.

## Discussion

Ferroptosis is caused by a redox imbalance between the production of oxidants and antioxidants, which is driven by the abnormal expression and activity of multiple redox-active enzymes that produce or detoxify free radicals and lipid oxidation products, whereas selective mitochondrial autophagy may inhibit ferroptotic death. Here, we show that reduced TXN leads to consequent reduced expression of BNIP3L, and reduced mitophagy, which ultimately causes a feedback loop that leads to increased ferroptosis (**Figure 7d**). Here, we found that TXN binds to the promoter of and activates the mitophagy-related protein BNIP3L, which was consistently down-regulated in aged GCs, blocking the occurrence of non-canonical mitophagy and resulting in the accumulation of damaged mitochondria and release of ROS in human GCs, in vitro cell models, and mouse models of aging.

Ferroptosis could be a potential therapeutic target for ART and infertility treatment. Notably, supplementation of NADH can rescue age-related declines in mouse fertility ^61,62^, possibly by inhibiting ferroptosis. Similarly, supplementation with antioxidants such as coenzyme Q10 could improve the quality of oocytes by reducing apoptosis in a mouse model ^63,64^, and human ART patients under 40 years old ^65^, although it remains controversial in clinical practice ^66^. In addition, ErZhiTianGui decoction (a mixture of ten herbs) was reported to regulate mitochondrial homeostasis, reduce ROS accumulation, and inhibit ACSL4-mediated lipid peroxidation in aged mice ^67^. Finally, direct inhibition of ferroptosis in mice by injection of deferoxamine improved fertility ^68^. Generally, antioxidant therapy by supplementation is thought to improve mitochondrial function, thereby improving oocyte quality, embryonic quality, and age-related reproductive outcomes. These data suggest ferroptosis could be both a marker of ovarian aging and a target for improved infertility ^68^. However, inhibition of ferroptosis in a mouse model of PCOS did not affect GC viability ^69^. Ultimately, the efficacy of direct intervention against ferroptosis in human patients remains unclear.

In conclusion, ovarian aging is closely associated with declining fertility and reproductive dysfunction, which has long been a challenging issue in reproductive medicine. We have determined that ferroptosis is associated with ovarian aging, and inhibiting ferroptosis could ameliorate some of the features of aging in a mouse model. We also discovered that TXN is down-regulated in aging GCs. TXN inhibits ferroptosis in aging GCs by binding (indirectly) to DNA and activating autophagy/mitophagy genes and BNIP3L-dependent mitophagy, therefore ameliorating ovarian aging. These results show how a TXN/BNIP3L-driven mitophagy pathway regulates ferroptosis in ovarian aging, thereby offering a basis for targeted prevention and delay of age-related ovarian dysfunction.

## Acknowledgments

This work was supported by grants from the National Natural Science Foundation of China (32270597 to A.P.H). The authors acknowledge the assistance of SUSTech Core Research Facilities.

## Author contributions

W.T.Y. designed the study. W.T.Y. and R.X. prepared figures, performed the experiments, analyzed the data, and wrote the first draft of the manuscript. Yi Liu performed experiments, analyzed the data, supervised the study, reviewed, and edited the manuscript. ZQ.H., Z.L. performed the bioinformatic analysis. M.G. analyzed the immunofluorescence. A.P.H. revised the manuscript, acquired funding, and assisted in the bioinformatic analysis. GQ.T. acquired funding and designed and supervised the study. All authors assisted in revising the manuscript, performing experiments, analyzing the data, and/or preparing figures.

## Ethical approval

The Institutional Review Board of the First Affiliated Hospital of Xi’an Jiaotong University approved this study (LLSBPJ-2024-250). All participants provided informed written consent. For animal studies, ethical approval was obtained from the Biomedical Ethical Committee of the Health Science Center of Xi’an Jiaotong University (XJTUAE2024-716). The use of animals in this research adheres to ethical guidelines, including minimizing pain and distress, providing proper housing and care, and using the minimum number of animals necessary to obtain valid scientific results.

## Data Accessions

The mass spectrometry data generated in this study were submitted to iProX ^70^ with the accession number: IPX0011098000. CUT&Tag sequence data was deposited in the Gene Expression Omnibus with accession number: GSE297159. This study includes a reanalysis of GSE255690 ^42^, GSE232306 ^4^, and GSE158802 ^60^.

