## Supplementary Material for "Thioredoxin-1 inhibits granulosa cell ferroptosis to rescue ovarian aging through mitophagy-dependent activation of BNIP3L"

#### Supplementary Tables

**Supplementary Table 1** – Table of unique ID, metabolite name, molecular weight, classes, positive/negative ion, mass spectrometry intensities, fold-changes, p-values, VIP status, and significantly up or down-regulated for young versus old human GCs, middle-aged versus young, and middle-aged versus old.

**Supplementary Table 2** – Table of Uniprot ID, gene description, gene name, mass spectrometry intensities, fold-changes, p-values, VIP status, and significantly up or down-regulated for young versus old human GCs, middle-aged versus young, and middle-aged versus old.

#### Materials and methods

##### Human sample collection

Human samples, including ovarian GCs and venous blood, were collected at the First Affiliated Hospital of Xi'an Jiaotong University from 2022 to 2023. Patients with diseases such as polycystic ovary syndrome (PCOS), premature ovarian insufficiency (POI), and other ovarian hypofunctions caused by tumor or endocrine diseases were excluded. Only patients requiring assisted reproductive treatment (ART) between 22-46 years old were included. Human ovarian GCs, follicular fluid, and blood samples were collected from 3 different age groups: old group (n=30, >36 years old), middle group (n=30, between 30-36 years old), and young group (n=30, ≤29 years old). Isolated cell purity was determined with 95% purity via follicle-stimulating hormone receptor staining. Peripheral blood was collected, and serum hormones were quantified using validated automated immunoassay methods (Elecsys electrochemiluminescence immunoassays on Cobas 6000, Roche Diagnostics).

The clinical characteristics of the patients were compared between the three age groups. BMI (23.94 ± 2.72 vs. 22.26 ± 2.67 vs 22.73 ± 4.13 $kg/m^2$, $P$ = 0.2173), infertility duration (4.08 ± 2.76 vs. 3.92 ± 1.98 vs. 2.52 ± 2.13 years, $P$ = 0.0982), menstrual cycle length (27.95 ± 1.12 vs 28.45 ± 2.13 vs. 28.14 ± 1.73 d, $P = 0.0623$), the basal E2, LH, progesterone and PRL all had no significant difference, while age (38.6 ± 2.03 vs 32.6 ± 1.68 vs 27.13 ± 1.97 years, $P < 0.001$), basal FSH (9.74 ± 4.12 vs. 7.12 ± 2.42 vs. 5.84 ± 2.87 mIU/mL, $P = 0.0024$), AMH (0.96 ± 1.03 vs. 2.06 ± 1.25 vs 4.27 ± 3.01 ng/mL, $P < 0.01$) and AFC (9.25 ± 3.45 vs 14.86 ± 4.32 vs. 18.45 ± 4.12, $P < 0.001$) had significant differences between the three groups.

**Mouse models**

Female C57BL/6 mice were provided by the Experimental Animal Center at Xi'an Jiaotong University. These mice were then raised on a 12-h light: 12-h dark cycle with *ad libitum* food and water. Mice were divided into a young group (12-week-old, n = 24) and an old group (52-week-old, n = 24). Treatment was given once a day for 3 weeks, using intraperitoneal injection containing 1mg/kg ferrostatin-1 (Fer-1) (MCE) for the old group, 20mg/kg erastin (MCE) for the young group, and sesame oil (Sigma-Aldrich) for the control group. Mice estrous cycles was determined by judging cell types of vaginal smears daily. Six mice from the vehicle (control) group and six mice from the experimental treatment group were euthanized to collect the bilateral ovaries and blood at the diestrus stage.

The remaining twelve mice were randomly divided into a control group (n = 6) and a TXN group (n = 6). Busulfan and cyclophosphamide (Aladdin) were dissolved in 0.9% saline containing 0.5% DMSO. Mice were treated with vehicle or busulfan (12 mg/kg) and cyclophosphamide (70 mg/kg) by intraperitoneal injection at 6:00 p.m. once a day (during this treatment, no longer given to mice) for 2 days. Also, all mice were euthanized to collect blood and bilateral ovaries.

For primary mouse GC isolation, ovaries were removed and punctured with 25-gauge needles (#5). The cells were pooled and filtered with a 40 μm cell strainer to remove oocytes. The isolated cells were determined to have 95% purity via follicle-stimulating hormone receptor staining. The mGCs (mouse GCs) were cultured in DMEM/F12 medium (Gibco BRL/Invitrogen) containing 10% fetal bovine serum

(HyClone), 1 mM sodium pyruvate, 2 mM glutamine, 100 IU/ml penicillin, and 100 μg/ml streptomycin.

##### Oocyte superovulation and *in vitro* fertilization in mice

C57BL/6 female mice were treated with peritoneal injections of 8 IU pregnant mare serum gonadotropin (PMSG) and followed 48~50 h later by 8 IU human chorionic gonadotropin (hCG) injections. Superovulated mice were sacrificed 12-14h after hCG injection, and the oviductal ampullae were broken to extrude the cumulus-oocyte complexes. An incubation period of approximately 2 h was sufficient for sperm capacitation. Approximately $2 \times 10^6$ sperm/ml were added to each IVF droplet. After 4-6 h incubation of oocytes and sperm, sperm and granulosa cells were gently removed by pipetting, and the number of ova with male and female pronuclei (2PNs) was counted. Sperm capacitation and IVF incubation were performed in 5% $CO_2$, 37°C, with saturated humidity. Pregnant mare serum gonadotropin (PMSG) and Human chorionic gonadotropin (hCG; Ningbo SANSHENG Biotechnology Co) were used.

##### Ovary hematoxylin and eosin (H&E) staining and follicle counting

Ovaries were fixed in 4% paraformaldehyde (pH 7.5) overnight at 4°C, dehydrated and embedded in paraffin, and cut into 5 μm slices, then stained with hematoxylin & eosin. The raw number of follicles was multiplied by five to determine the total number of follicles, as the sampling fraction was every 5th section being counted. The follicles were classified according to the classification system used in a previous study [1]. Both ovaries from three mice of each group were used for the analysis.

##### Human granulosa cell isolation and culture

After oocyte retrieval from patients, the human cumulus oophorus was digested with 0.1% hyaluronidase (Fujifilm Irvine Scientific) for 45-60 seconds. The collected cells were digested into single-cell suspensions using 0.25% trypsin-EDTA (Gibco) and then separated by centrifugation at 500 g in a swing bucket centrifuge for 5 min. After isolation, incubation of human GCs was conducted in DMEM/F12 containing 10% FBS and 1% (v/v) penicillin/streptomycin (Sigma-Aldrich) as previously reported.

Human steroidogenic GC-like tumor cell line (KGN) was obtained from Procell and cultured with DMEM/F12 containing 10% (v/v) FBS and 1% (v/v) penicillin-streptomycin at 37°C in an atmosphere of 5% $CO_2$. KGN cells were treated with these concentrations of $H_2O_2$ (400 μM), Fer-1 (5 μM), Erastin (10 μM), PX-12 (5 μM), TXNIP-IN-1 (10 μM), and NADPH (100 μM), unless otherwise indicated.

**Immunohistochemistry and Immunofluorescence**

Mice ovaries were fixed in buffered 4% (w/v) paraformaldehyde, embedded in paraffin, sectioned to approximately 5 μm, and put on glass slides. The ovarian sections were deparaffinized in xylene, rehydrated, and retrieved by microwave heating with buffer of Citrate Antigen Retrieval Solution (Beyotime, P0081) for 20 min. Endogenous peroxidase activity was quenched by incubation with 3% $H_2O_2$ for 10 min. After blocking in 3% BSA for 30 mins, tissues were stained with primary antibodies at 4°C overnight as followed: rabbit anti-human/mouse FSHR (1:200, WLA032, Wanlei), rabbit anti-human/mouse $p21^{CDNK1A}$ (1:200, WL0362, Wanlei), mouse anti-human/mouse LMNB1 (1:200, sc-374015, Santa Cruz), rabbit anti-human/mouse TXN (1:200, 14999-1-AP, Proteintech), rabbit anti-human/mouse SLC7A11 (1:400, XJ3706481, Invitrogen), mouse anti-human/mouse GPX4 (1:200, sc-166570, Santa Cruz) and rabbit anti-human/mouse NOX4 (1:200, 14347-1-AP, Proteintech). The sections were subsequently incubated with an anti-mouse or rabbit biotinylated secondary antibody with an HRP-polymer antibody at RT for 30 min (PV-8000, Zhongshan Goldenbridge), stained with a DAB mixture (ZLI-9018, Zhongshan Goldenbridge) for 3-5 min, and observed under a microscope.

For Immunofluorescence, cells were fixed in 4% (w/v) paraformaldehyde for 30 min at room temperature and then permeated with 1% Triton X-100 at room temperature for 20 min. Then cells were incubated at 4 °C overnight with primary antibody: rabbit anti-human/mouse FSHR (1:100, WLA032, Wanlei), mouse anti-human/mouse TXN (1:100, sc-166393, Santa Cruz), rabbit anti-human/mouse BNIP3L

(1:100, 12986-1-AP, Proteintech), rabbit anti-human/mouse TXN (1:100, 14999-1-AP, Proteintech), rabbit anti-human/mouse SLC7A11 (1:200, XJ3706481, Invitrogen), mouse anti-human/mouse GPX4 (1:100, sc-166570, Santa Cruz), and rabbit anti-human/mouse NOX4 (1:200, 14347-1-AP, Proteintech). After washing with PBS 3 times thoroughly, GCs were incubated at 37 °C for 1 h with fluorescence-conjugated secondary antibody: goat anti-mouse IgG (DyLight 488, 1:400, RS23210, Immunoway) and goat anti-rabbit IgG (AbFluor 555, 1:400, RS3411, Immunoway). The cell nucleus was stained with 4′6-diamidino-2-phenylindole (DAPI, Vector Laboratories) and then viewed using a fluorescence microscope (Ti2-E, Nikon).

**Senescence-associated beta-galactosidase (SA-β-Gal) staining**

To stain for SA-β-Gal, fixed cells were given the staining solution mix as recommended by the manufacturer (Servicebio). Staining was performed overnight at 37°C in an incubator without $CO_2$ control. After the overnight incubation, cells were washed twice with PBS to stop the staining.

**Cell viability (CCK8 assay)**

Cells were seeded in 96-well plates at a density of 2~5×$10^3$cells/well and cultured for 24 h. Then, cells were treated with the indicated agents at various concentrations as per the experimental design. Cell viability was assessed using the Cell Counting Kit-8 (CCK-8) assay (CT001, SparkJade). Briefly, at the end of the treatment time, 10 μL of CCK-8 solution was added to each well, and the plates were incubated for an additional 1~4 h. The absorbance at 450 nm was measured using a microplate reader. The cell viability was calculated as a percentage of the absorbance of the treated cells relative to the untreated control cells. The data were collected from at least three independent experiments, and each experiment was performed in triplicate to ensure reproducibility.

**Western blot**

Total protein extraction was conducted as previously reported. The total of 30μg protein was resolved and then transferred to PVDF membranes. Membranes were blocked with

5% fat-free milk for 1 h at room temperature and immunoblotted with primary antibodies at 4°C overnight. The antibodies used in the present study included: mouse anti-human LMNB1 (1:500, sc-374015, Santa Cruz), rabbit anti-human p21$^{CDNK1A}$ (1:500, WL0362, Wanlei), rabbit anti-human SLC7A11 (1:800, XJ3706481, Invitrogen), mouse anti-human GPX4 (1:500, sc-166570, Santa Cruz), rabbit anti-human TXN (1:500, 14999-1-AP, Proteintech), rabbit anti-human NOX4 (1:500, 14347-1-AP, Proteintech), rabbit anti-human TXNRD1 (1:500, 11117-1-AP, Proteintech), mouse anti-human NRF2 (1:500, sc-365949, Santa Cruz), mouse anti-human YAP (1:500, sc-376830, Santa Cruz), mouse anti-human BCL2 (1:500, sc-7382, Santa Cruz), mouse anti-human ACSL4 (1:500, sc-365230, Santa Cruz), rabbit anti-human/mouse BNIP3L (1:500, 12986-1-AP, Proteintech), rabbit anti-human LC3I/II (1:500, WL01506, Wanlei), rabbit anti-human p53 (1:500, 60283-2-Ig, Proteintech), mouse anti-human p16$^{CDNK2A}$ (1:500, sc-1661, Santa Cruz) and mouse anti-human GAPDH (1:8000, YM3029, Immunoway). The PVDF membranes were subsequently incubated with HRP-conjugated goat anti-mouse or goat anti-rabbit secondary antibodies at a dilution of 1:5000 (RS0083 and RS0076, Immunoway), and enhanced chemiluminescent substrate (ECL, WBULP-100ML, Millipore, Merck) was used for protein blot detection. Signals were visualized using an imaging system (Tanon-5200), and ImageJ software (version 1.54g) was used to measure protein expression.

**Malondialdehyde (MDA), Glutathione (GSH), and ROS measurement**

MDA was measured using the MDA Content Assay Kit (Solarbio) following the manufacturer's instructions. Briefly, the cell extracting solution or follicular fluid (100 μL) was incubated with the test solution (300 μL) at 100°C for 60 min and centrifuged at 10000g for 10 min. The OD value of the supernatants was measured at 532 and 600 nm and used to calculate the MDA content.

Cells were harvested by scraping, and glutathione was measured using the Glutathione Assay Kit (Solarbio) according to the manufacturer's protocol. GSH concentrations were calculated using a standard curve and normalized to the total protein level in each sample. Three independent biological replicates were performed for each condition.

ROS was measured using 2′,7′-dichlorofluorescein diacetate ($H_2DCF$-DA, D399, Invitrogen-ThermoFisher) by flow cytometry. KGN cells were treated with $H_2O_2$ and ferroptosis inducer or inhibitor for 24 h at 37 °C, and then they were incubated in DMEM/F12 medium containing 5 μM $H_2DCF$-DA for 30 min at room temperature. Fluorescence intensity was determined using a flow cytometer (Agilent Novocyte).

**BODIPY 581/591 C11 analysis**

The day before the experiment, $2\times10^5$ cells per well were seeded into 6-well dishes. The treatment medium was removed, and cells were washed twice with HBSS, then labeled with 5 μM BODIPY 581/591 C11 (D3861, Invitrogen-ThermoFisher) and incubated at 37°C for 20 min. The label mixture was removed, and 1 mL of fresh HBSS was added to the cells. The BODIPY 581/591 C11 value was calculated as the ratio of green fluorescence (oxidized probe) to total (green + red, which indicates total reduced plus oxidized probe) fluorescence by assessing the percentage of oxidized probe by flow cytometry (FACS, Agilent NovoCyte) and analyzed using NovoExpress. The cellular compartmentalization of the oxidized C11-BODIPY 581/591 was monitored by tracking green fluorescence with a fluorescence microscope (Ti2-E, Nikon).

**Measurement of mitochondrial membrane potential**

Mitochondrial membrane potential assay kit with JC-1 (CBIC2, HY-15534, MCE) was purchased to evaluate the mitochondrial membrane potentials ($\Delta\varphi m$). $2\times10^5$ cells were performed in 6-well plates and incubated with a working solution containing 10 μM JC-1 in a 5% $CO_2$ atmosphere at 37 °C for 20 min. The fluorescence intensity of the JC-1 monomers/aggregates (green fluorescence for monomer, red fluorescence for aggregate) was measured using fluorescence microscopy. The calculation results of the green/red fluorescence ratio were used to assess mitochondrial membrane potential. The same samples were used to evaluate the mitochondrial membrane potential by assessing the percentage of monomers and aggregates by flow cytometry and analyzed using NovoExpress.

**Real-time PCR (RT-PCR)**

Total RNA was extracted using an RNA isolation kit (G3640-50T, Servicebio) following the manufacturer's instructions. Samples were reverse transcribed using the SweScript RT II First Strand cDNA Synthesis Kit (G3332-50, Servicebio). RNA concentrations were measured using a Nanodrop 2000 Spectrophotometer (Biolab) at a wavelength of 260 nm. Real-time PCR assays were performed with a Bio-Rad CFX96 real-time PCR instrument (Bio-Rad) with three technical replicates for all assays. The relative expression of genes was calculated with the comparative threshold cycle (CT) method as $2^{-\Delta\Delta CT}$. The primers used include: *TXN*-F: GTGAAGCAGATCGAGAGCAAG, *TXN*-R: CGTGGCTGAGAAGTCAACTACTA; *GPX4*-F: GAGGCAAGACCGAAGTAAACTAC, *GPX4*-R: CCGAACTGGTTACACGGGAA; *SCL7A11*-F: TCTCCAAAGGAGGTTACCTGC, *SLC7A11*-R: AGACTCCCCTCAGTAAAGTGAC; *ELOVL5*-F: TAACAGGAGTATGGGAAGGCA, *ELOVL5*-R: ACCAGAGGACACGGATAATCTT; *NOX4*-F: CAGATGTTGGGGCTAGGATTG, *NOX4*-R: GAGTGTTCGGCACATGGGTA; *NNMT*-F: ATATTCTGCCTAGACGGTGTGA, *NNMT*-R: TCAGTGACGACGATCTCCTTAAA; *GAPDH*-F: CTGGGCTACACTGAGCACC, *GAPDH*-R: AAGTGGTCGTTGAGGGCAATG.

**Mitophagy Autophagy detection in GCs.**

This study employed a GPP-RFP-LC3 dual-fluorescence viral transfection system (Genomeditech, Shanghai) to monitor autophagic activity. Cells were plated at $2\times10^5$ cells/well in 12-well plates the day before transfection or treatment, and then transduced with the GPP-RFP-LC3 reporter system using lentiviral/adenoviral vectors with optimized multiplicity of infection (MOI) following cell seeding. The system leverages

GPP (green fluorescent protein) to label nascent autophagosomes and RFP (red fluorescent protein) to track lysosomal degradation, followed by confocal microscopy to quantify autophagic flux via the number of RFP-GFP double-positive puncta (yellow dots). GCs were also stained with the Mitophagy Detection Kit (Dojindo) for mitophagy, according to the manufacturer's instructions.

**Transmission electron microscopy (TEM)**

1 $mm^3$ tissue blocks from ovaries were transferred into fresh TEM fixative at room temperature with 3% glutaraldehyde and 1% osmium acid for 2 hours. After dehydration with isoamyl acetate, samples were dried and treated with vacuum spraying. Prepared samples were examined under TEM, and images were collected using an HT7800 transmission electron microscope (Hitachi).

**Cell transfection**

Cell transfection was performed as previously described. Briefly, cells cultured in 6-well plates were transfected with 2 μg *TXN*-cDNA or sh*TXN* plasmid for overexpression or knockdown of the expression of TXN, and a shRNA targeting LUC (luciferase) or a scrambled RNA sequence acted as a negative control. The *TXN* cDNA (NM_003329.4) was amplified and inserted into the pEX-3 (pGCMV/MSC/Neo) vector backbone. For the sh*TXN* plasmid, the target sequences of AGGTGATAAACTTGTAGTAGTTG and ACCATTAATGAATTAGTCTAATC were cloned into the vector pGPU6/Neo. Also, the sh*BNIP3L* plasmid was constructed with the target of GTCAGAAGAAGAAGTTGTAGAAG with pGPU6/Neo vector backbone. All the vectors were synthesized and were transfected using Lipofectamine 2000 Transfection Reagent (Invitrogen) according to the manufacturer’s protocol.

**Dual-luciferase reporter assay**

The *BINP3L*-promoter LUC reporter plasmids were built containing the sequence of the *BINP3L*-promoter region from -3452 to -2600 and transiently co-transfected into KGN cells ($5\times10^4$) plated in a 24-well plate dish with pTK-RL plasmid. The activity of

Luciferase and Renilla was measured 48 h post-transfection using the Dual Luciferase Assay kit (Promega) according to the manufacturer's instructions. All experiments were performed in triplicate. The efficiency of transfection was normalized to the paired Renilla luciferase activity, and the specific activity was displayed as the fold change of the experimental group versus the control group.

**Quantitative chromatin immunoprecipitation**

Quantitative chromatin immunoprecipitation (qChIP) assays were performed according to the manufacturer's protocol for the EZ-ChIP Assay Kit (Cat. #17–371). Chromatin-protein complexes were immunoprecipitated with 10 μg of anti-TXN antibodies and 20 μL of fully resuspended protein A/G magnetic beads. For the negative control, 1 μg of normal mouse IgG was used. Real-time PCR was performed to amplify the regions of interest or internal negative control regions. Each sample was assayed in triplicate, and the fold enrichment ratio was calculated as the value of the ChIP sample versus the corresponding input sample. Samples that yielded a twofold enrichment or better were considered positive targets. The primers used for these studies are listed: Region 1 (R1)-F: CTCCTAACAAAGTCTTCGA, Region 1 (R1)-R: GTCAGGGGCTGAGACTGAAG; Region 2 (R2)-F: ACAAATTGTCCGCCTT, Region 2 (R2)-R: CTAATACTAACGAATAC.

**Bulk and single-cell RNA-seq data analysis**

Bulk RNA-seq data were aligned to the genome using the STAR aligner as previously described [2]. Reads were mapped to genes using the hg38 genome assembly and GENCODE v42 transcriptome using STAR[3], and mapped to genes using te_count/scTE [2]. Data was normalized using EDASeq [4]. Differential expression analysis of two conditions/groups was performed using DESeq2 (1.20.0) [5]. The resulting p-values were adjusted using Benjamini-Hochberg correction. Genes with an adjusted $p\text{-value} < 0.05$ and $\log_2 FC > 1$ were considered differentially expressed. Downstream analysis was performed using glbase [6].

Single-cell RNA-seq reads were aligned to the human genome using STAR-solo as previously described [2], genes were quantified using te_counts/scTE [2], and downstream analysis was performed using scanpy [7].

**CUT&Tag of TXN**

CUT&Tag was performed as previously described [8], with some modifications. CUT&Tag assay was performed using the NovoNGS CUT&Tag 3.0 High-Sensitivity Kit (N259-YH01-01B, Novoprotein) according to the manufacturer's instructions. Briefly, around 100,000 cells for each sample were used to perform CUT&Tag with a primary antibody against TXN (14999-1-AP, Proteintech).

Analysis was performed as previously described [9]. Briefly, cutadapt was used to filter reads: -a CTGTCTCTTATACACATCTCCGAGCCCACGAGAC -A CTGTCTCTTATACACATCTGACGCTGCCGACGA --quality-cutoff 10 -m 50. Reads were aligned to the hg38 genome using bowtie2 with the settings -I 10 -X 1000 [10], and peaks were detected using MACS2 with default settings and a q-value of 0.01 [11]. TFBS motif discovery was performed using HOMER using findMotifsGenome.pl with default arguments [12]. CistromeGO was used to annotate KEGG pathways or GOBP gene sets to TXN binding[13]. Other analysis was performed using glbase3 [6].

**Mass spec non-targeted metabolism and protein data analysis.**

DIA quantitative proteomic detection was performed using a . Take out the sample from the -80°C refrigerator, transfer it to a 1.5ml centrifuge tube, and add an appropriate amount of DB protein solution (8 M urea 100 mM TEAB, pH 8.5), Shake and mix well, and sonicate in an ice water bath for 5 minutes to fully lyse. Centrifuge at 4°C and 12000 g. After 15 minutes, 1M DTT was added, and allowed to react at 56°C for 1 hour. Ice bath for 2 minutes, then add sufficient iodoacetamide at room temperature in the dark for 1 hour.

Take protein samples and add DB protein solution (8 M urea, 100 mM TEAB, pH 8.5) to make up the volume. Add 100 μL of trypsin and 100 mM TEAB buffer, mix well, and perform enzyme digestion at 37°C for 4 hours. Then add trypsin and $CaCl_2$. Enzyme digest overnight. Add formic acid to adjust the pH to less than 3, mix well, centrifuge at room temperature at 12000 g for 5 minutes, and slowly pass the supernatant through C18 desalination column, followed by continuous cleaning with cleaning solution (0.1% formic acid, 3% acetonitrile) 3 times, and then adding an appropriate amount of eluent (0.1% Formic acid, 70% acetonitrile), collect the filtrate, and freeze dry.

Prepare mobile phase A (100% water, 0.1% formic acid) and B (80% acetonitrile, 0.1% formic acid). Use 10 μL, dissolve freeze-dried powder in solution A, centrifuge at 14000 g for 20 minutes at 4°C, take 200ng of the supernatant sample for injection, and perform liquid quality testing. Send Upgrade the UHPLC system with Vanquish Neo, with a C18 pre column size of 174500 (5mm × 300 μm, 5 μm), heating in a 50°C column incubator, the C18 analytical column is ES906 (PepMap TM Neo UHPLC 150 μm x 15 cm, 2 μm; Thermo Fisher).

Spectrometer, Easy spray (ESI) ion source, set ion spray voltage to 1.9kV, and ion transfer tube temperature to 290°C, The mass spectrometry adopts a data dependent acquisition mode, with a full scan range of m/z 380-980 for the primary mass spectrometry, and a resolution set for the primary mass spectrometry for 240000 (200 m/z), the AGC is set to 500%, the size of the mother ion window is set to 2-Th, and the number of DIA windows is 300, NCE set to 25%, secondary m/z acquisition range is 150 to 2000.

Search and analyze raw files using the DIA-NN search software based on the Uniprot (homo_sapiens_uniprot_2023_3_13.fasta) protein database. The search parameters are set as follows: The mass tolerance of precursor ions is 10 ppm, and the mass tolerance of fragment ions is 0.02 Da. Fixed modification to cysteine alkylation modification, variable modification to methionine oxidation modification, N-terminal modification to acetylation modification, methionine loss, and methionine loss+acetylation, allowing up to one cleavage site to be missed. To improve the quality of the analysis results, DIA-NN software further filters the retrieval results: spectra with a credibility of over 99% peptide Spectrum Matches (PSMs) are trustworthy PSMs that only retain reliable spectral peptides and proteins. Perform FDR validation to remove peptides and proteins with FDR greater than 1% FC<1 [fold change, FC] is defined as a differentially expressed protein (DEP).

Non-targeted metabolism experiments were performed using a Q Exactive HF/Q Exactive HF-X (Thermo Fisher) and a Vanquish UHPLC (Thermo Fisher). Metabolite extraction was performed as described: Briefly. Cells were placed in a tube. Add 300 μL of 80% methanol aqueous solution. Freeze in liquid nitrogen for 5 minutes; Vortex for 30 seconds after melting on ice and sonicate for 6 minutes. Centrifuge at 5000 rpm and 4 ℃ for 1 minute, transfer the supernatant to a new centrifuge tube, and freeze dry it into a dry powder. Dissolve the corresponding 10% methanol solution according to the volume of the sample taken and inject it into the LC-MS for analysis. Blank sample: 53% methanol aqueous solution is used as a substitute for the experimental sample, and the pre-treatment process is the same as that of the experimental sample. Instrument Parameters were set to: Chromatographic conditions
Chromatographic column: Hypersil Gold column (C18) (Thermo Fisher), Column temperature: 40°C, Flow rate: 0.2 mL/min, Positive mode: Mobile phase A: 0.1% formic acid, Mobile phase B: Methanol, Negative mode: Mobile phase A: 5 mM ammonium acetate, pH 9.0. Mobile phase B: Methanol. Mass spectrometry conditions: Select the scanning range of m/z 100-1500; The settings of the ESI source are as follows: spray voltage: 3.5 kV; sheath gas flow rate: 35 psi; auxiliary gas flow rate flow rate: 10 L/min; ionic transport tube temperature: 320°C; Ion introduction into radio frequency electricity S-lens RF level: 60; Aux gas heater temp: 350°C, polarity: positive, negative.

Import the downloaded data (. raw) file into CD 3.3 search software for processing, and perform analysis on each metabolite, simple screening of retention time, mass to charge ratio, and other parameters, followed by peak area correction using the first QC to enable identification. Then set the quality deviation to 5 ppm, signal strength deviation to 30%, minimum signal strength, and extract peaks based on information such as ions, quantify peak areas, integrate target ions, and then separate them, molecular formula prediction of sub-ion peaks and fragment ions using mzCloud.

Compare mzVault and Masslist databases, remove background ions using blank samples, and quantify the original data. The results are based on the formula: original quantitative value of the sample/(sum of quantitative values of metabolites in the sample/QC1 quantitative value of metabolites in the sample), normalize the sum to obtain the relative peak area, and calculate the CV of the relative peak area in the QC sample.

The statistical significance of each metabolite between the groups was based on a t-test, p-value was determined, and the fold change of metabolites between the groups was calculated. A metabolite was considered statistically different if VIP>1, p-value<0.05, and FC⩾2 or FC⩽0.5. Other analysis was performed using glbase [6].

**Statistical analysis**

Statistical analysis was performed with SPSS version 18.0 (SPSS Inc.). Measurement data were analyzed with mean ± standard deviation; The two-tailed $\chi^2$ test or Fisher's exact test was used to determine the significance of the differences between covariates. A univariate analysis was performed using Student's *t*-test (two tailed) or the Mann-Whitney U-test. For comparison among groups, the $\chi^2$ test or one-way ANOVA was performed. A p-value < 0.05 was considered statistically significant, unless otherwise indicated.

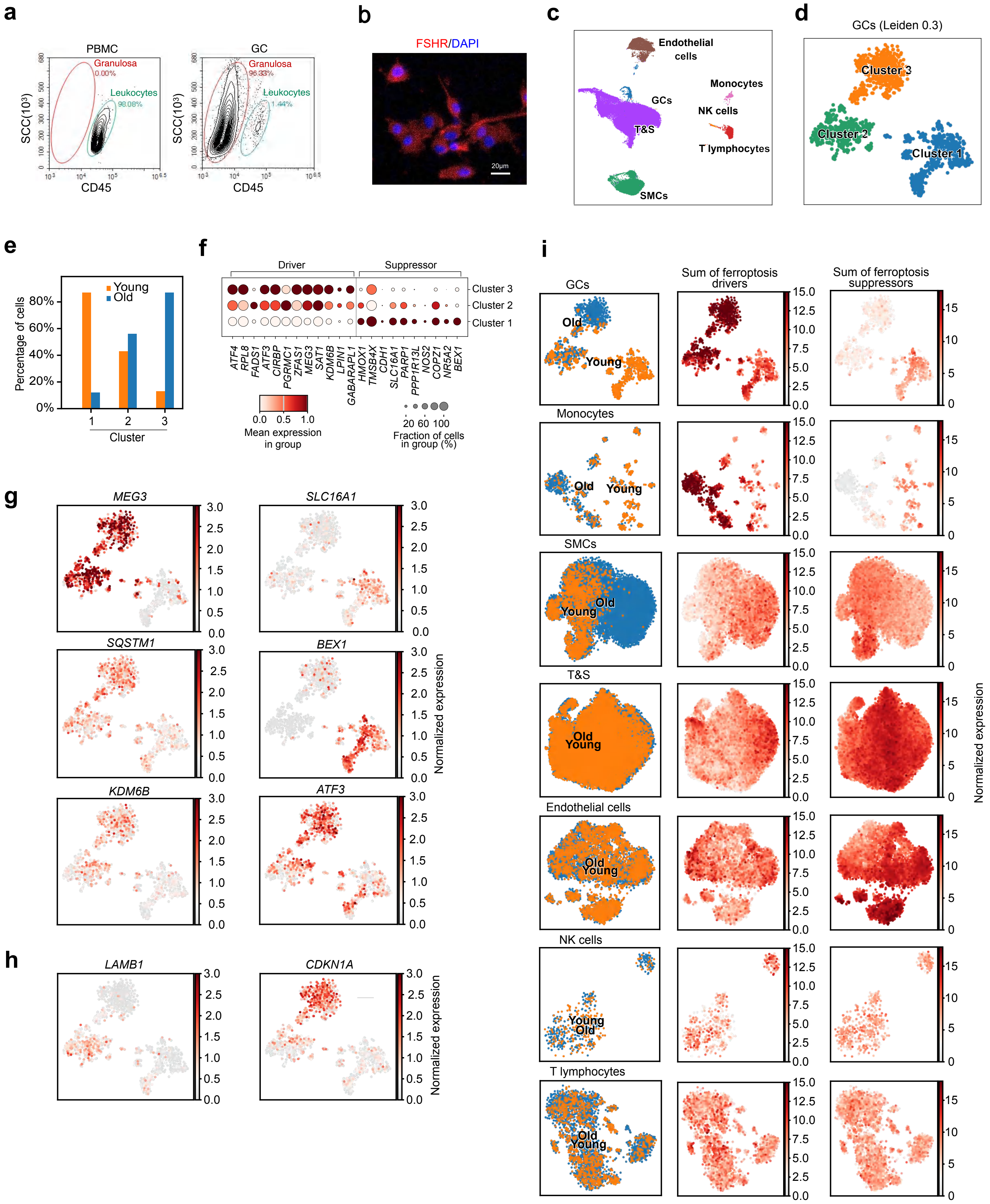


Supplementary Figure 1

**Supplementary Figure 1. Purification of GCs from human and single-cell RNA-seq analysis of ferroptosis.**

**a** Flow cytometry of CD45 levels in GCs (right). CD45+ levels in peripheral blood mononuclear cells (PBMCs) were used as a negative control (left).

**b** Fluorescence images of GCs stained with FSHR.

**c** UMAP (uniform manifold and projection) of all ovary cells. GCs = granulosa cells; T&S = Theca and Stromal cells; NK cells = Natural Killer cells; SMCs = smooth muscle cells. Data is from a reanalysis of Ref. [114], for this panel and panels D-I.

**d** tSNE (t-stochastic neighbor embedding) of the GCs alone. Clusters were determined using the Leiden algorithm with a resolution of 0.3.

**e** Bar chart of the percentage of cells from the young or old group in each cluster.

**f** Bubble plot of selected ferroptosis drivers and suppressors in the three clusters defined in **panel d**.

**g** tSNE of GCs colored by the normalized expression of the indicated genes.

**h** tSNE colored by the expression of the indicated cell cycle (*CDKN1A*) or nuclear lamina genes (*LAMB1*).

**i** tSNE plots of the indicated cell types (left-most column) and the sum of expression of ferroptosis-driver and suppressor genes. T&S = Theca and Stromal cells; NK cells = Natural Killer cells; SMCs = smooth muscle cells. Data is from a reanalysis of Ref. [114].

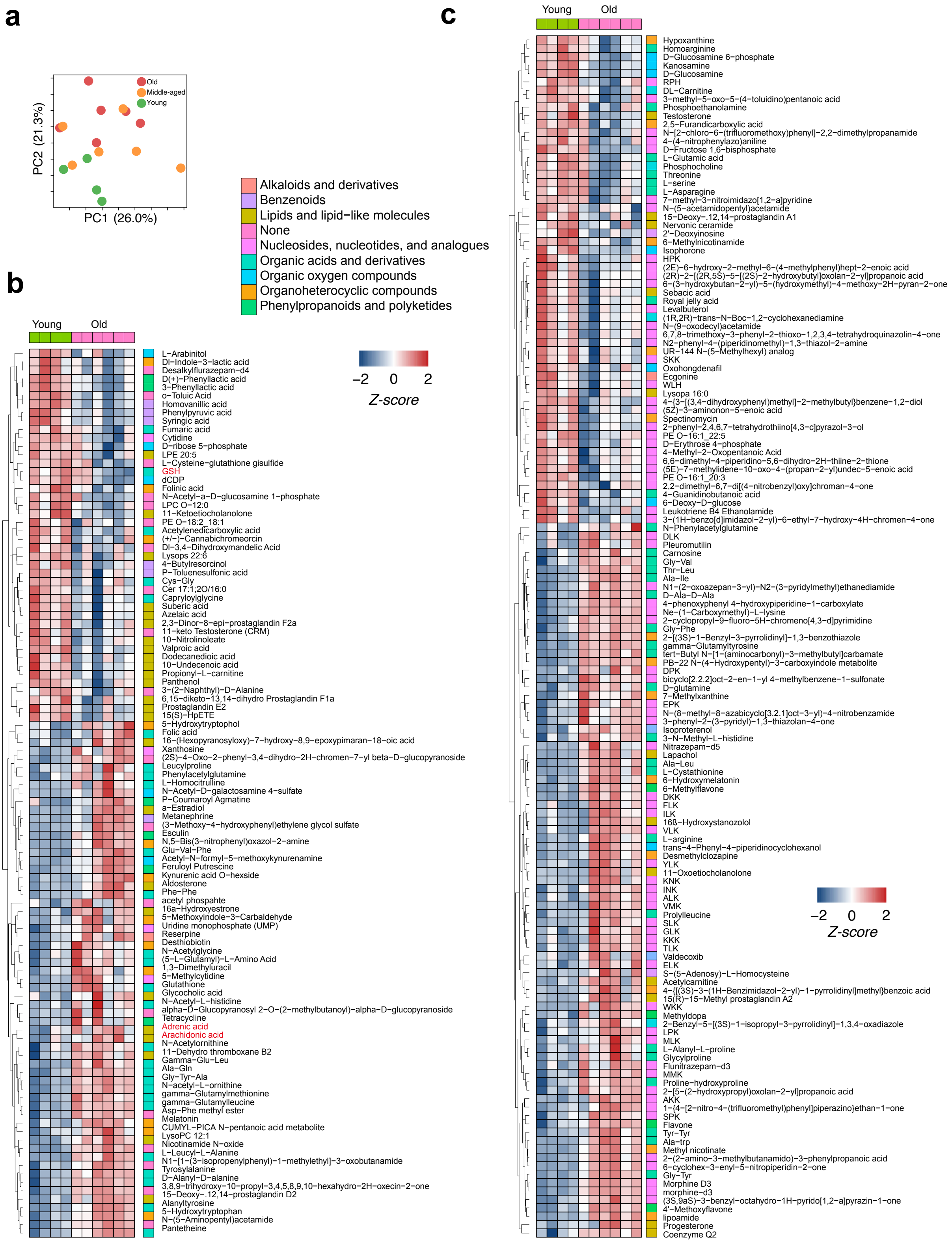


### Supplementary Figure 2

**Supplementary Figure 2. Non-targeted metabolomics of GCs from old, middle and young women.**

**a** PCA (principal component analysis) of the metabolic mass spec results for old, middle-aged, and young GCs from human ovaries.

**b** Heatmap of the Z-scores for significantly different negative ion metabolites in young and old human GCs.

**c** Heatmap of the Z-scores for significantly different positive ion metabolites in young and old human GCs.

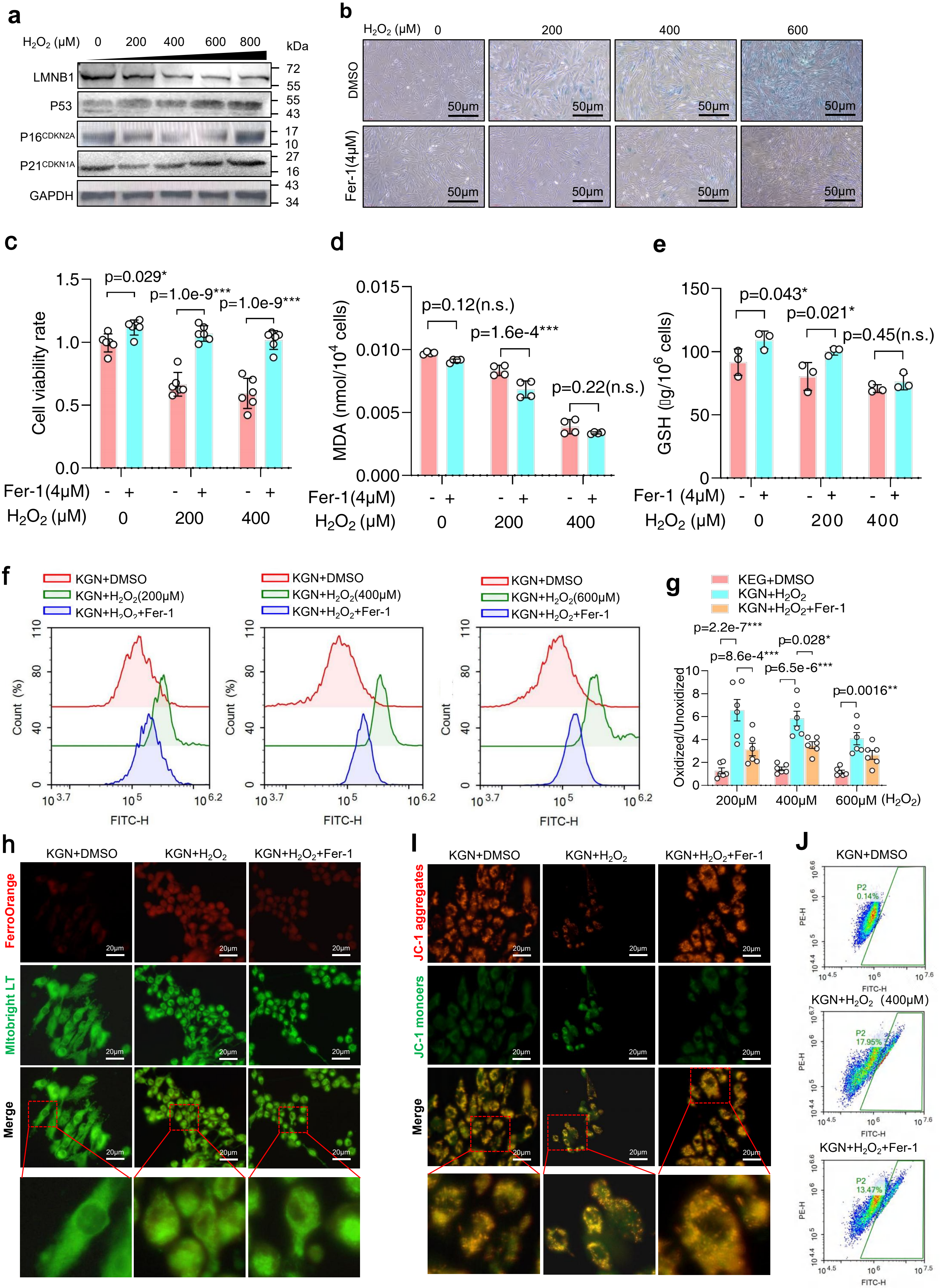


**Supplementary Figure 3**

**Supplementary Figure 3. Increased ferroptosis in $H_2O_2$-treated KGN cells**

**a** Western blot for LMNB1, p53, $p16^{CDNK2A,}$ and $p21^{CDNK1A}$ in KGN cells treated with increasing concentrations of $H_2O_2$.

**b** SA-β-gal staining in KGN cells treated with increasing concentrations of $H_2O_2$ and DMSO (control) or Fer-1.

**c** Bar chart representing cell viability rate by CCK8 assay for KGN cells treated with $H_2O_2$ and DMSO or Fer-1. Significance is from an independent two-sided Student's *t* test.

**d** Bar chart of MDA levels in KGN cells treated with increasing concentrations of $H_2O_2$. Significance is from an independent two-sided Student's *t* test.

**e** Bar chart of GSH levels in KGN cells treated with increasing concentrations of $H_2O_2$. Significance is from a two-sided Student's *t-test*.

**f** Example flow cytometry histograms for oxidized lipids in KGN cells treated with increasing levels of $H_2O_2$ and Fer-1 based on C11-BODIPY 581/591 dye.

**g** Bar chart quantifying the ratio of oxidized to unoxidized lipids (as measured by C11-BIODIPY) in KGN cells treated with DMSO (vehicle), $H_2O_{2,}$ or H2O2 + Fer-1. Statistical significance was determined through a one-way ANOVA test, followed by Tukey's test for pairwise comparisons between groups. The experiment was performed six times.

**h** Staining of KGN cells with the indicated treatments for FerroOrange ($Fe^{2+}$; red) and mitochondria (Mitobright LT; green).

**i** Staining of KGN cells with the indicated treatments for JC-1 monomers (green) and aggregates (red).

**j** Flow cytometry plot for JC-1 monomers (displayed in green) and aggregates (displayed in red) in KGN cells that received the indicated treatments.

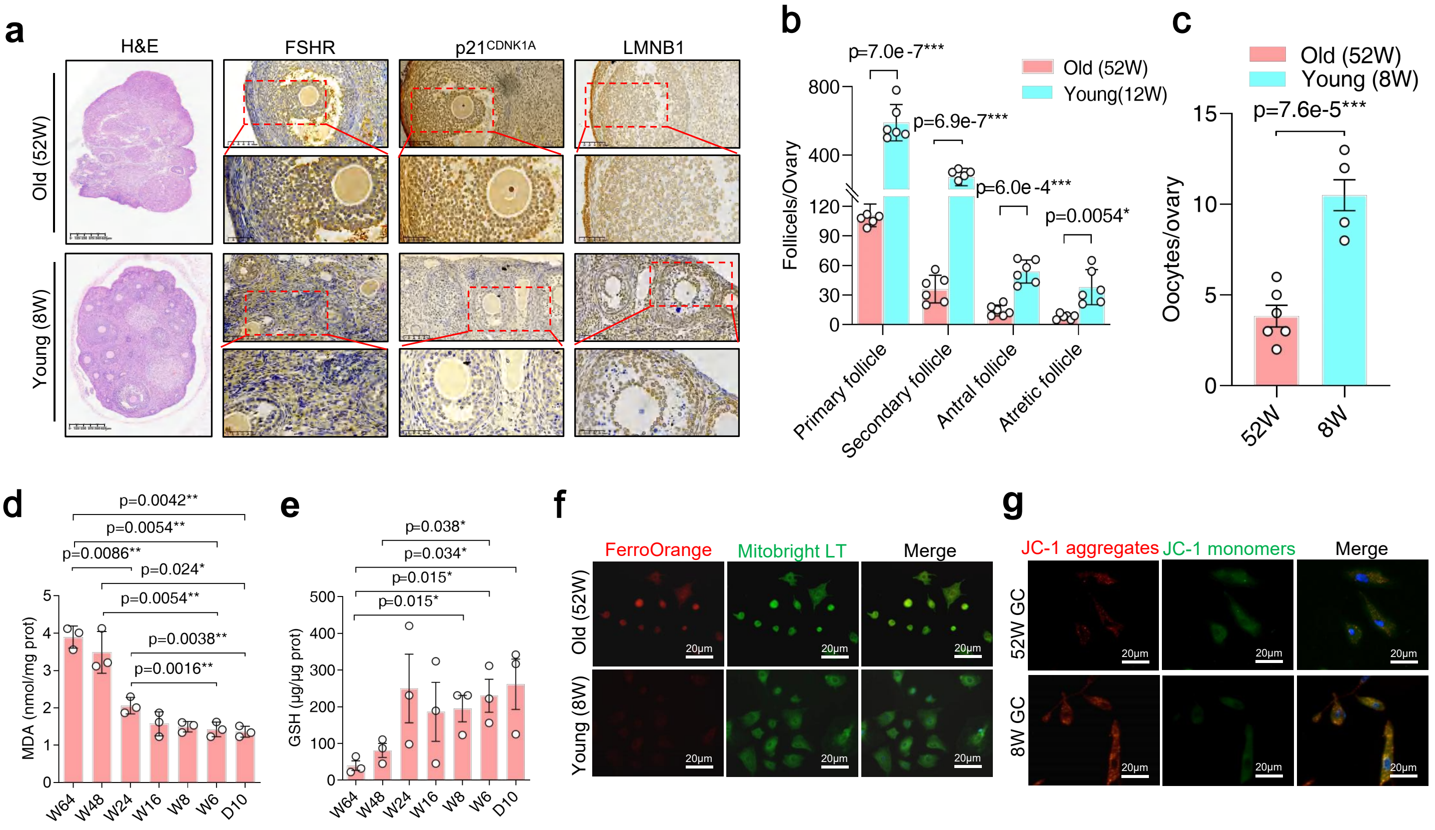


### Supplementary Figure 4

**Supplementary Figure 4. Increased markers for ferroptosis in the ovaries of old mice**

**a** Hematoxylin and eosin staining and IHC of FSHR, $p21^{CDKN1A,}$ and LMNB1 in mouse ovaries at 52 weeks and 8 weeks old (n=3).

**b** Dot plot showing the number of primary, secondary, antral, and atretic follicles of mice (n=6) at 52 weeks old and 8 weeks old. Significance is from a two-sided Student's *t-test*.

**c** Dot plot showing the number of oocytes per ovary slice in young and old mice (n=3). Significance is from a two-sided Student's *t-test*.

**d** Dot plots of MDA in mice from young (Week 6/W6) to old (Week 64/W64) and postnatal (Day 10/D10). Statistical significance was determined through a one-way ANOVA test, followed by Tukey's test for pairwise comparisons between groups.

**e** Dot plots of GSH in mice from young (Week 6/W6) to old (Week 64/W64) and postnatal (Day 10/D10). Statistical significance was determined through a one-way ANOVA test, followed by Tukey's test for pairwise comparisons between groups.

**f** Fluorescence microscopy of GCs from old and young mouse ovaries stained with FerroOrange (Fe2+) and Mitobright LT. Scale bar = 20 μm.

**g** Fluorescence microscopy of GCs from old and young mouse ovaries stained with JC-1. Scale bar = 20 μm.

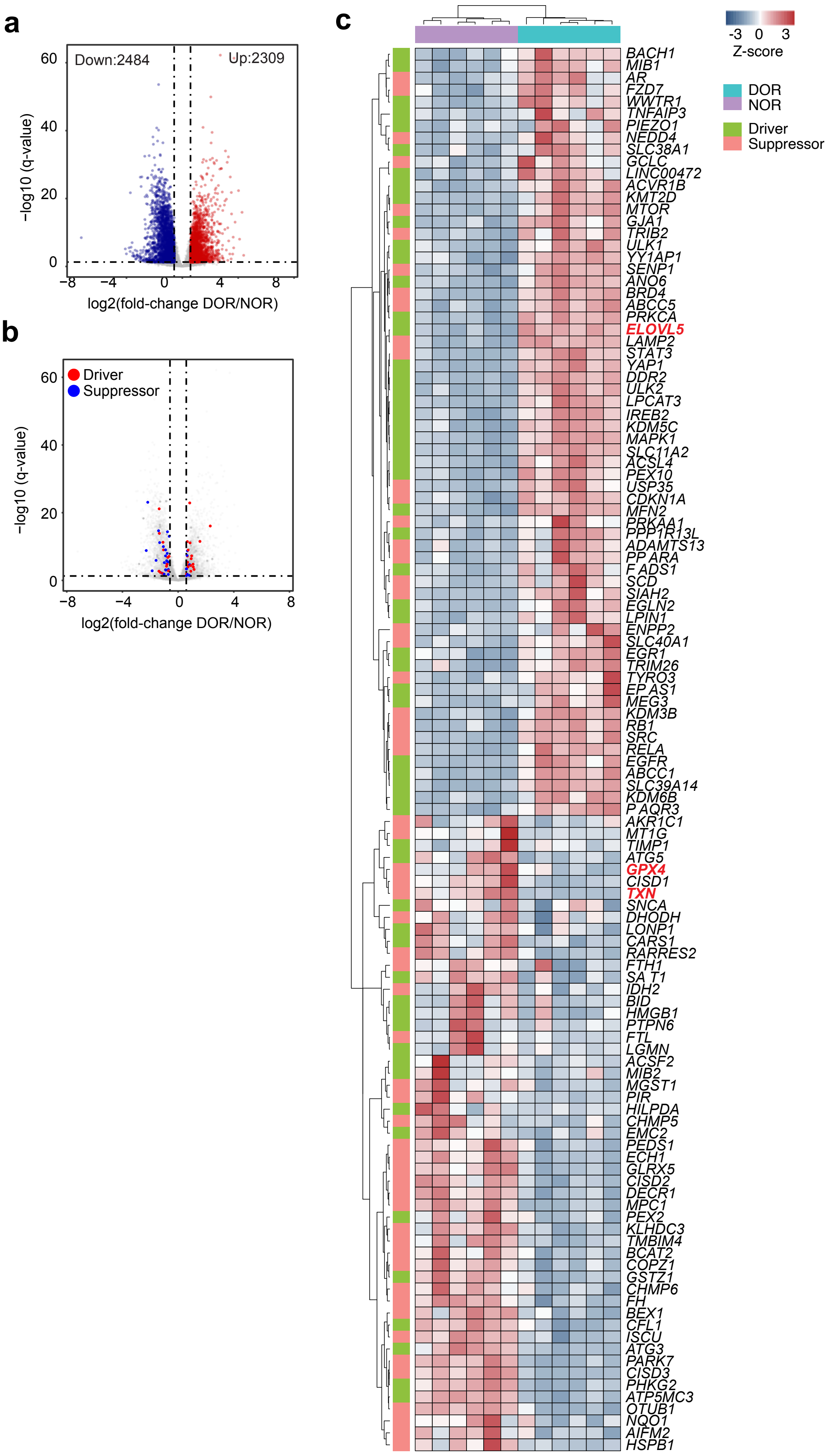


**Supplementary Figure 5**

**Supplementary Figure 5. Ferroptosis-related genes are elevated in depleted ovarian reserve GCs**

**a** Volcano plot of all genes significantly differentially expressed genes for NOR (normal ovarian reserve) (n=6) versus DOR (depleted ovarian reserve) (n=6) patients. A gene was considered significantly differentially expressed if its absolute fold-change was > 2.0 and the Bonferroni-Hochberg corrected p-value (q-value) was <0.01. Data is from a reanalysis of GSE232306 [2][15].

**b** As in **panel a**, but labelling only ferroptosis driver or suppressor genes, as defined in FerrDB[16].

**c** Heatmap of all significantly differentially regulated (as defined in **panel a**) drivers and suppressors (as defined in panel B) of ferroptosis in DOR and NOR patients.

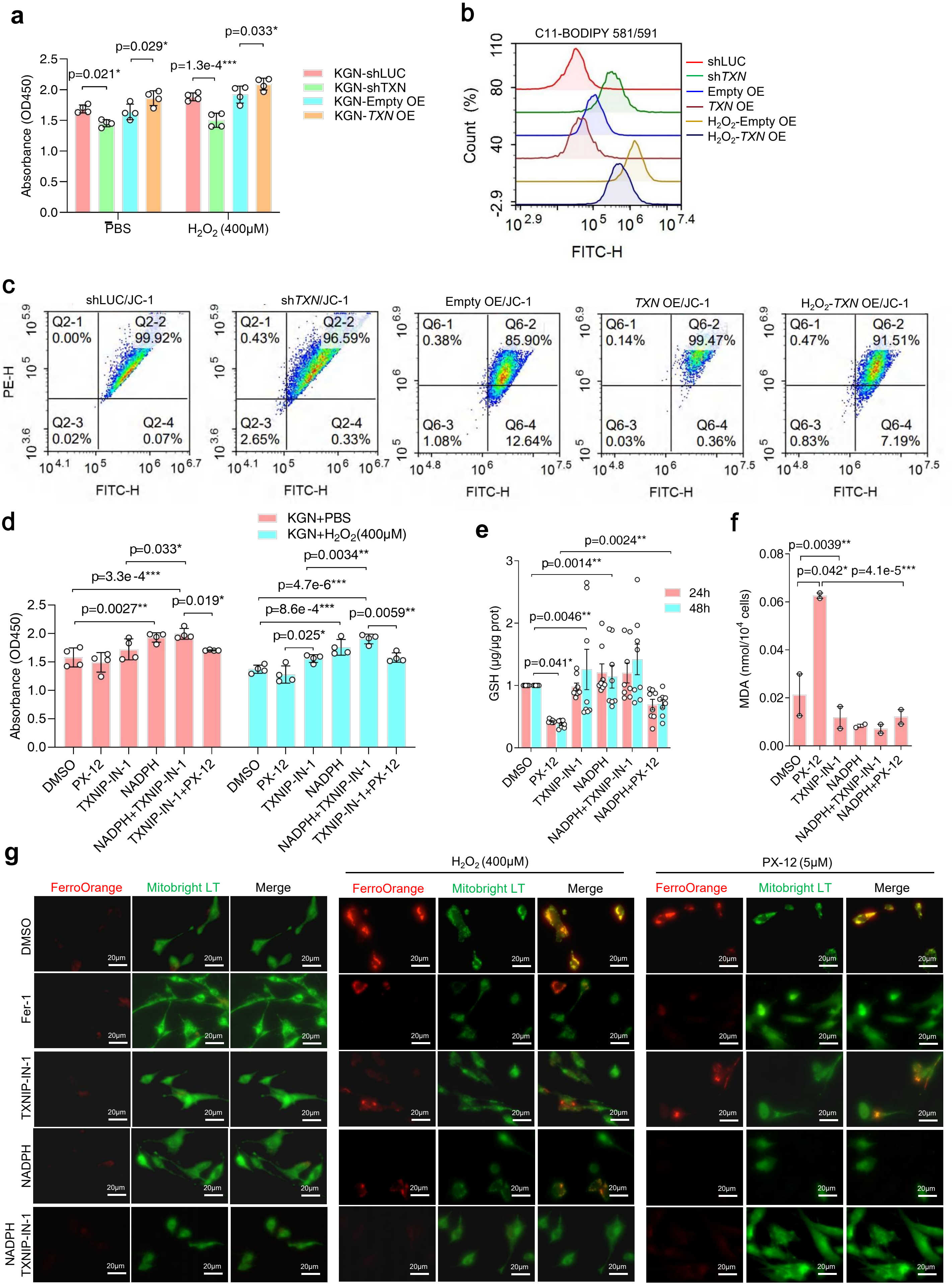


**Supplementary Figure 6**

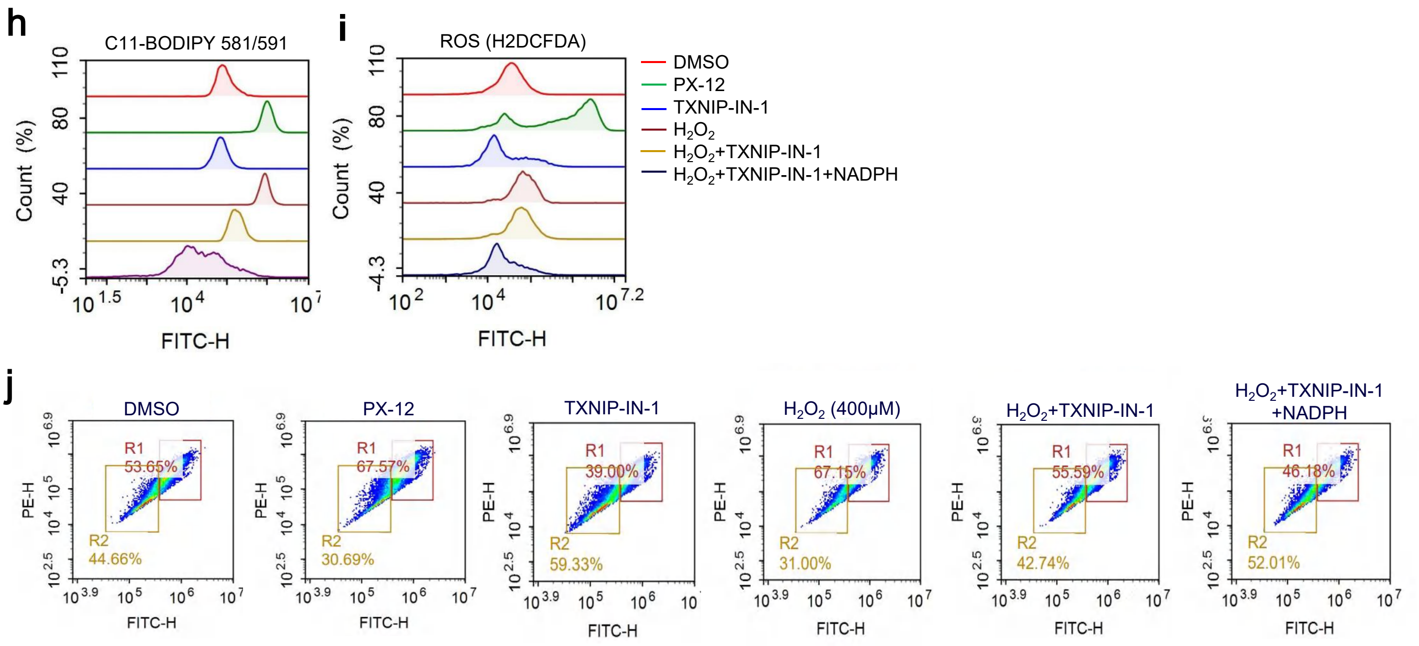


### Supplementary Figure 6

**Supplementary Figure 6. TXN modulates the ferroptosis response**

**a** Bar chart showing the cell viability of KGN cells transfected with shRNAs targeting *LUC* (control) or *TXN*, or an Empty overexpression vector, or a vector containing *TXN* (*TXN* OE). The cells were treated with $H_2O_2$ or PBS (Control). Statistical significance was determined through a one-way ANOVA test for overall group comparison, followed by Tukey's test for pairwise comparisons between groups.

**b** Example flow cytometry histograms of C11-BODIPY stained oxidized lipids in KGN cells transfected with an shRNA targeting *TXN* or overexpressing *TXN* (*TXN* OE) in cells treated with $H_2O_2$ or PBS.

**c** Example flow cytometry histograms of JC-1 levels (mitochondrial membrane potential) in KGN cells transfected with an shRNA targeting *TXN* or *LUC* as a control, or an Empty vector, or an overexpression vector containing *TXN*. The cells were also treated with $H_2O_2$ or PBS.

**d** Dot plots showing the cell viability for KGN cells treated with the TXN inhibitor (PX-12) or activators (TXNIP-IN-1 and NADPH) alone or in combination. Significance is from a one-way ANOVA test, followed by Tukey's test for pairwise comparisons between groups.

**e** Dot plot of GSH levels in KGN cells after treatment with a TXN inhibitor (PX-12) or activators (TXNIP-IN-1 and NADPH) alone or in combination. Significance is from a one-way ANOVA test for overall group comparison, followed by Tukey's test for pairwise comparisons between groups.

**f** Dot plot of MDA levels in KGN cells after treatment with the TXN inhibitor (PX-12) or activators (TXNIP-IN-1 and NADPH) alone or in combination. Significance is from a one-way ANOVA test for overall group comparison, followed by Tukey's test for pairwise comparisons between groups.

**g** Fluorescence images of KGN cells treated with Fer-1, TXN-IN-1, NADPH, $H_2O_2$, or the TXN inhibitor PX-12. $Fe^{2+}$ accumulation was measured using the FerroOrange fluorescent probe (red) and mitochondria by MitoBright LT (green).

**h** Example flow cytometry histograms of C11-BODIPY stained oxidized lipids in KGN cells treated with the TXN inhibitor (PX-12) or activators (TXNIP-IN-1 and NADPH) alone or in combination and with $H_2O_2$.

**i** Example flow cytometry histograms of ROS as measured by DCFDA-stained oxidized lipids in KGN cells treated with the TXN inhibitor (PX-12) or activators (TXNIP-IN-1 and NADPH) alone or in combination and with $H_2O_2$.

**j** Example flow cytometry plots of mitochondrial membrane potential analyzed by JC-1 monomers (displayed in green) and aggregates (displayed in red) in KGN cells that received the indicated treatments.

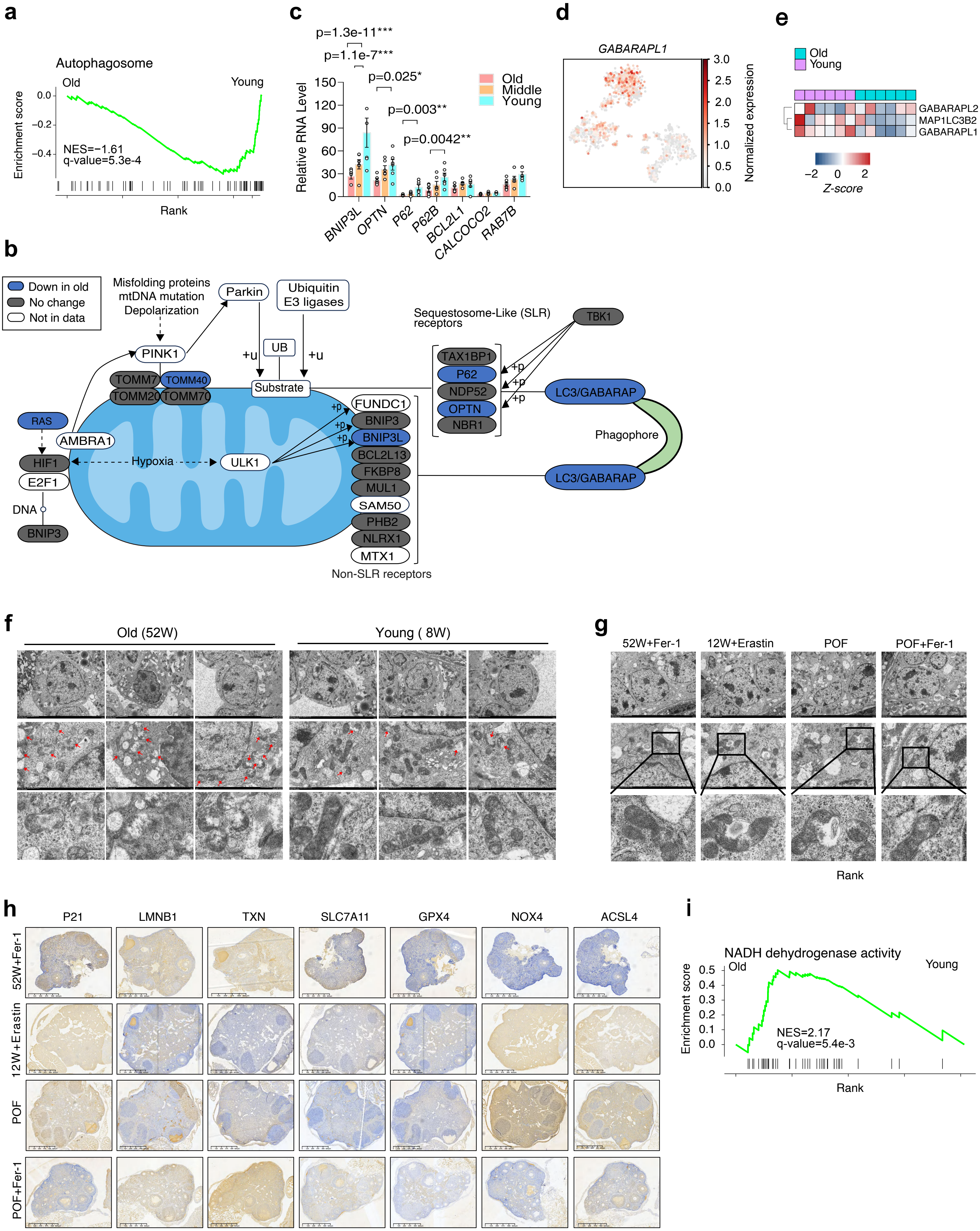


Supplementary Figure 7

**Supplementary Figure 7. Mitochondrial function is dysregulated in old GCs in humans and mice**

- **a** GSEA of protein mass spectrometry from young and old human GCs, ranked by their fold-change from old to young. NES=normalized enrichment score, q-value=Bonferroni-Hochberg corrected p-value.
- **b** Schematic of proteins involved in mitophagy. Down-regulated proteins in the GC human mass spectrometry data are marked in blue, unchanged in grey, and proteins not detected in the mass spec in white. Adapted from the mitophagy figure from KEGG hsa04137.
- **c** Dot plot showing the RT-qPCR results for the indicated genes in old, middle-aged, and young human GCs. Significance is from a one-way ANOVA test for overall group comparison, followed by Tukey's test for pairwise comparisons between groups.
- **d** tSNE of GCs colored by the normalized expression of the LC3-component GABARAPL1. Data is from a reanalysis of Ref. [114].
- **e** Heatmap of the selected LC3-related proteins detected from mass spectrometry for the LC3 proteins GABARAPL2, GABARAPL1, and MAP1LC3B2.
- **f** Mitochondrial ultrastructure detected by TEM in ovarian tissues from young mice (8 weeks old, n = 3) and old mice (52 weeks old, n = 3).
- **g** Mitochondrial ultrastructure detected by TEM in ovarian tissues from young mice (8 weeks old, n=3) and old mice (52 weeks old, n=3) treated with a Ferroptosis inhibitor (Fer-1) activator (Erastin) or in POF model mice treated with Fer-1.
- **h** Immunohistochemistry of FSHR, $p21^{CDKN1A}$, LMNB1, TXN, SLC7A11, GPX4, NOX4, and ACSL4 in mouse ovaries at 52 weeks, and treated with Fer-1 and 12 weeks old treated with Erastin, and in POF mice and POF mice treated with Fer-1 (n=3).
- **i** GSEA for young and old human GCs, showing significant enrichment of NADH dehydrogenase activity in old human GCs.

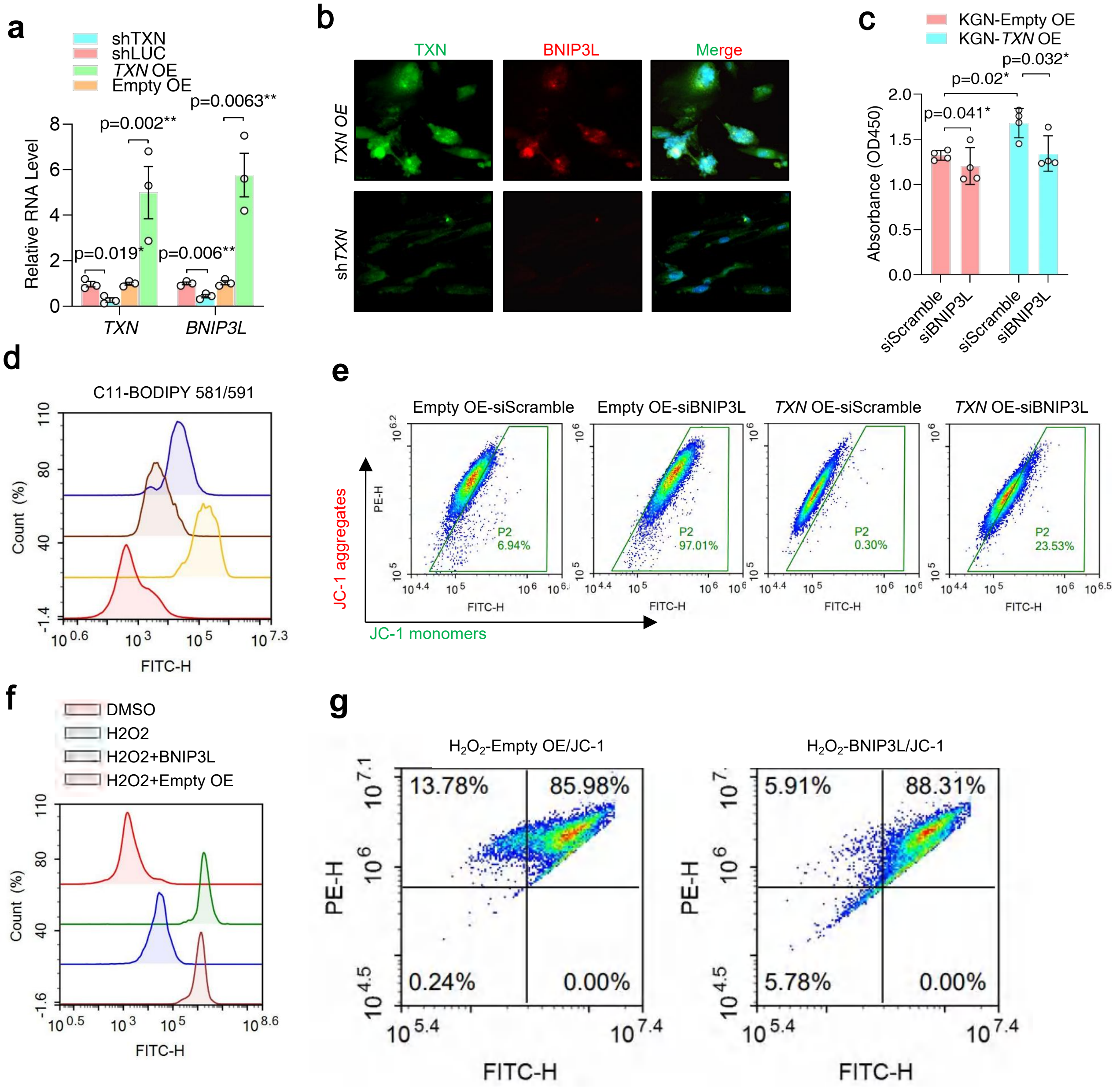


**Supplementary Figure 8**

**Supplementary Figure 8. TXN regulates BNIP3L**

**a** Dot plot showing RT-qPCR data for *TXN* and *BNIP3L* in KGN cells transfected with shRNAs targeting *LUC* (Control), *TXN*, or an empty overexpression vector or one containing *TXN*. Significance is from a one-way ANOVA test, followed by Tukey's test for pairwise comparisons between groups.

**b** Immunofluorescence microscopy images of GCs stained with TXN or BNIP3L in *TXN* OE cells or *TXN* knockdown KGN cells.

**c** Gene ontology for BP (biological process) pathways for TXN-bound genes, generated using CistromeGO.

**d** Genome view of the *SLC7A11* locus.

**e** TXN binding sites (-2600-3000bp) in the promoter regions of BNIP3L were predicted with the database of Promoter 2.0 Prediction Server.

**f** ChIP-qPCR for TXN bound to the *BNIP3L* promoter region in *TXN* overexpressing KGN cells. Significance is from a two-way ANOVA with Bonferroni correction.

**g** Cell viability (CCK8 assay, measured by absorbance OD450) in KGN cells transfected with shRNAs targeting *BNIP3L* or a scramble shRNA (control) with an overexpression vector containing *TXN* or an empty vector,

**h** GSH and MDA levels in KGN cells transfected with either an empty overexpression vector or a vector containing TXN, and in combination with an shRNA targeting BNIP3L, or a scrambled sequence as a control. Significance is from a two-way ANOVA with a Bonferroni test.

**i** Example flow cytometry histograms of C11-BODIPY stained oxidized lipids in KGN cells treated with the TXN inhibitor (PX-12) or activators (TXNIP-IN-1 and NADPH) alone or in combination and with $H_2O_2$.

**j** Example flow cytometry plots of JC-1 aggregates (Red) and monomers (green) in KGN cells transfected with an shRNA targeting *BNIP3L* or *LUC* as a control, or an Empty vector, or an overexpression vector containing *TXN* (*TXN* OE).

**k** Example flow cytometry histograms of C11-BODIPY stained oxidized lipids in KGN cells treated with the TXN inhibitor (PX-12) or activators (TXNIP-IN-1 and NADPH) alone or in combination and with $H_2O_2$.

**l** Example flow cytometry plots of JC-1 aggregates (Red) and monomers (green) in KGN cells transfected with an shRNA targeting *BNIP3L* or *LUC* as a control, or an Empty vector, or an overexpression vector containing *TXN* (*TXN* OE). The cells were also treated with $H_2O_2$.

**Fig 1b**

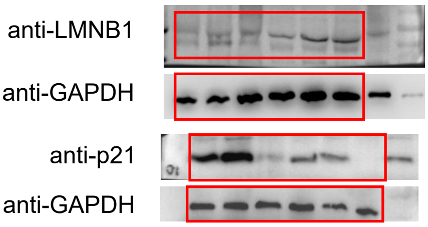


**Fig 3e**

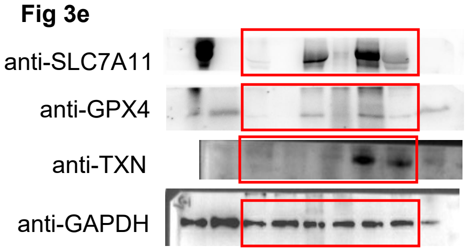


**Fig 4a-up**

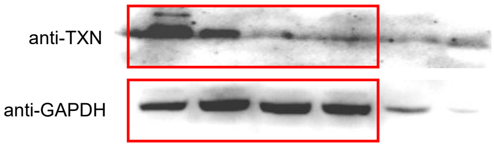


**Fig 4a-down**

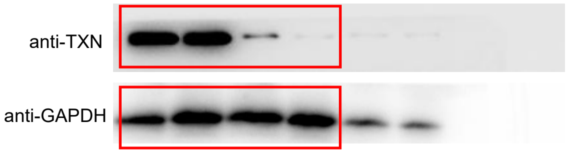


**Fig 4e-left**

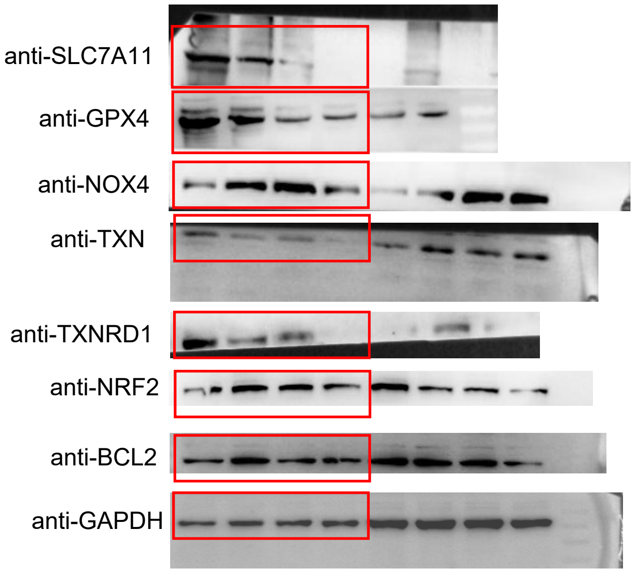


**Fig 4e-right**

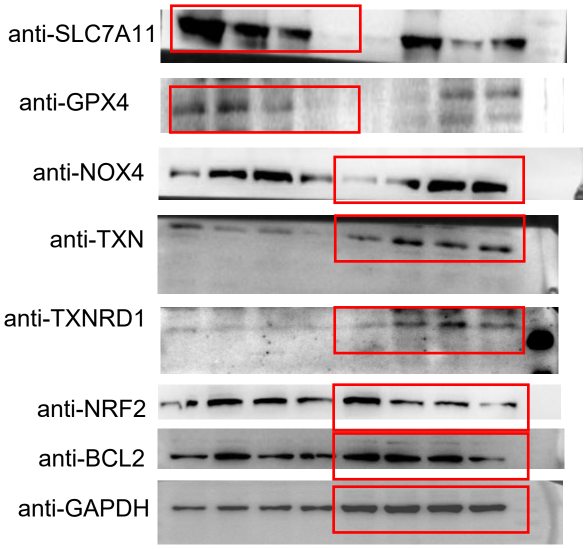


**Fig 4g**

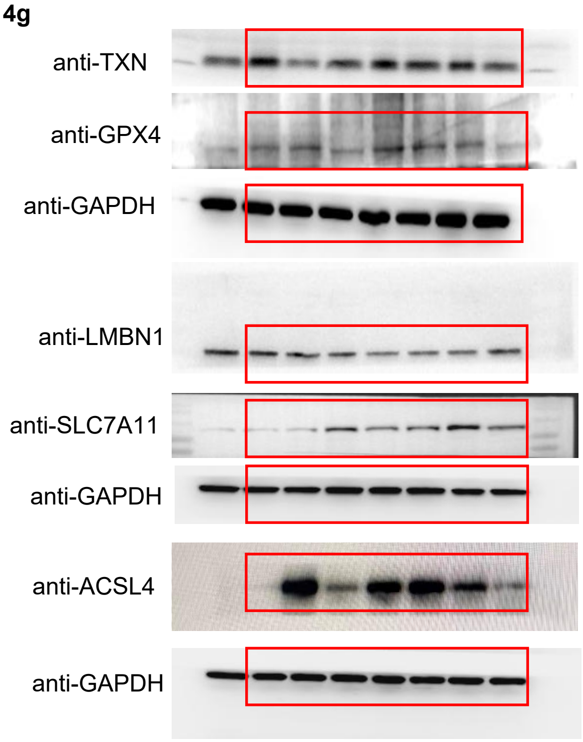


**Fig 5d**

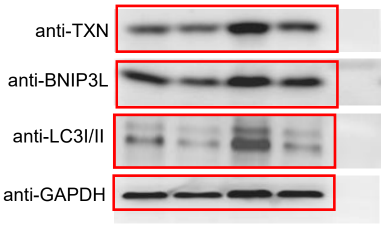

**Fig 6b**

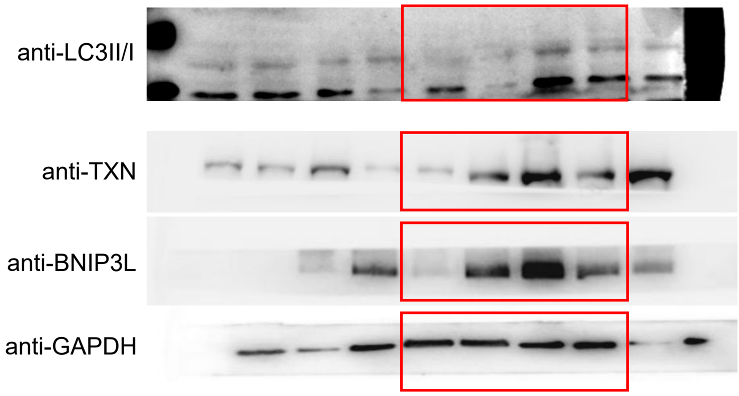


**Fig 6c**

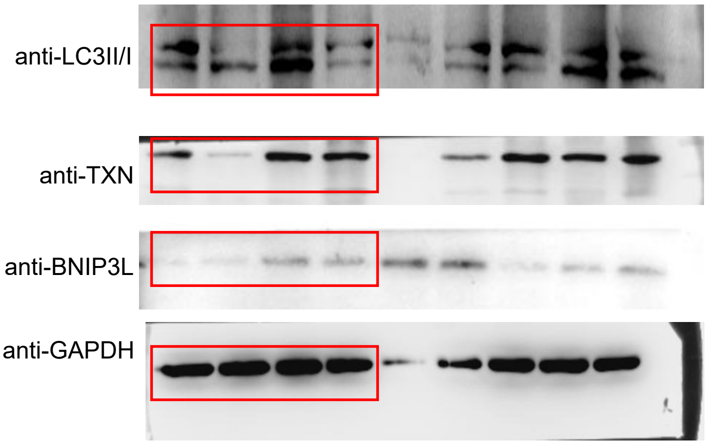


**Fig 6l**

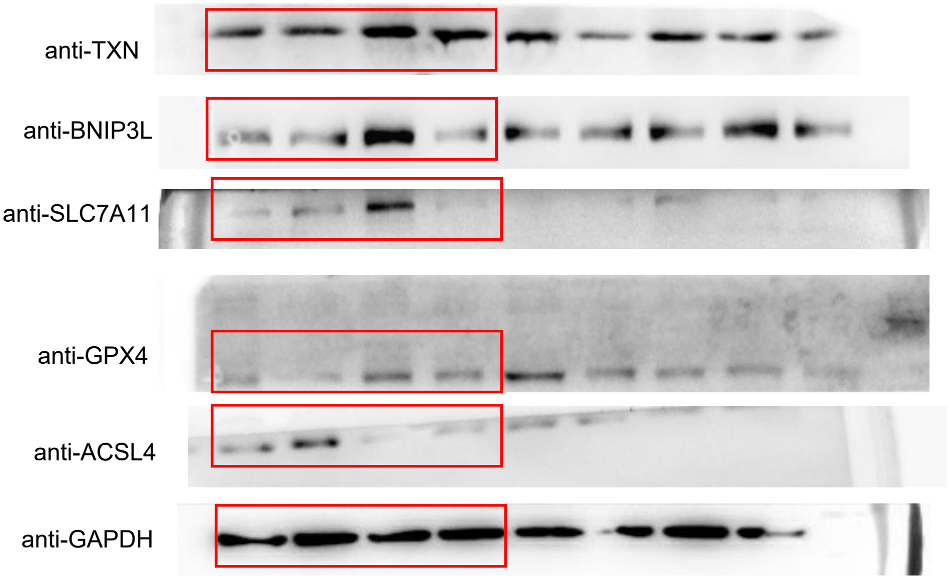


**SFig 3a**

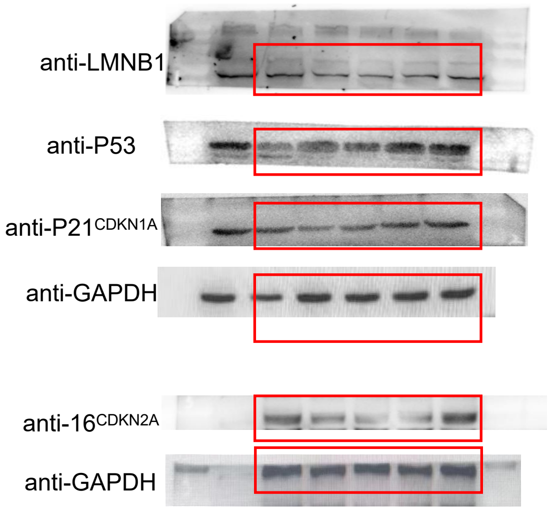
